# Establishment of a pancreatic adenocarcinoma molecular gradient (PAMG) that predicts the clinical outcome of pancreatic cancer

**DOI:** 10.1101/2020.03.25.998138

**Authors:** Rémy Nicolle, Yuna Blum, Pauline Duconseil, Charles Vanbrugghe, Nicolas Brandone, Flora Poizat, Julie Roques, Martin Bigonnet, Odile Gayet, Marion Rubis, Samir Dou, Nabila Elarouci, Lucile Armenoult, Mira Ayadi, Aurélien de Reyniès, Marc Giovannini, Philippe Grandval, Stephane Garcia, Cindy Canivet, Jérôme Cros, Barbara Bournet, Louis Buscail, BACAP Consortium, Vincent Moutardier, Marine Gilabert, Juan Iovanna, Nelson Dusetti

## Abstract

**BACKGROUND:** A significant gap in pancreatic ductal adenocarcinoma (PDAC) patient’s care is the lack of molecular parameters characterizing tumors and allowing a personalized treatment. The goal of this study was to examine whole PDAC transcriptomic profiles to define a signature that would predict aggressiveness and treatment responsiveness better than done until now.

**METHODS AND PATIENTS:** Tumors were obtained from 76 consecutive resectable (n=40) or unresectable (n=36) tumors. PDAC were transplanted in mice to produce patient-drived xenografts (PDX). PDX were classified according to their histology into five groups, from highly undifferentiated to well differentiated. This classification resulted strongly associated with tumors aggressiveness. A PDAC molecular gradient (PAMG) was constructed from PDX transcriptomes recapitulating the five histological groups along a continuous gradient. The prognostic and predictive value for PMAG was evaluated in: i/ two independent series (n=598) of resected tumors; ii/ 60 advanced tumors obtained by diagnostic EUS-guided biopsy needle flushing and iii/ on 28 biopsies from mFOLFIRINOX treated metastatic tumors.

**RESULTS:** A unique transcriptomic signature (PAGM) was generated with significant and independent prognostic value. PAMG significantly improves the characterization of PDAC heterogeneity compared to non-overlapping classifications as validated in 4 independent series of tumors (e.g. 308 consecutive resected PDAC, HR=0.321 95% CI [0.207;0.5] and 60 locally-advanced or metastatic PDAC, HR=0.308 95% CI [0.113;0.836]). The PAMG signature is also associated with progression under mFOLFIRINOX treatment (Pearson correlation to tumor response: -0.67, p-value < 0.001).

**CONCLUSION:** We identified a transcriptomic signature (PAMG) that, unlike all other stratification schemas already proposed, classifies PDAC along a continuous gradient. It can be performed on formalin-fixed paraffin-embedded samples and EUS-guided biopsies showing a strong prognostic value and predicting mFOLFIRINOX responsiveness. We think that PAMG could unify all PDAC preexisting classifications inducing a shift in the actual paradigm of binary classifications towards a better characterization in a gradient.

**Trial Registration:** The PaCaOmics study is registered at www.clinicaltrials.gov with registration number NCT01692873. The validation BACAP study is registered at www.clinicaltrials.gov with registration number NCT02818829.

## Introduction

Pancreatic ductal adenocarcinoma (PDAC) is one of the most aggressive gastrointestinal tumors. While activating mutations in *KRAS* are the most common genetic alterations [1], mutations in other driver genes such as *CDKN2A, TP53* or *SMAD4* are randomly associated to *KRAS* mutations, generating a heterogeneous genetic landscape between patients. However, these mutations do not predict patient outcome or tumor drug sensitivity and PDAC patients with similar clinical presentation show high variability in overall survival (OS), ranging from 3 months to >5-6 years after diagnosis. While histopathological analyses of tumors revealed OS is shorter in patients presenting with aggressive poorly-differentiated tumors relative to patients with well-differentiated ones [2], this analysis required large amounts of undamaged tumor tissue. Such samples are only available from resected tumors, representing as few as 15% of PDAC cases. For resectable PDAC, the current recommendation is upfront surgical resection followed by systemic chemotherapy with or without radiation [3]. However, this strategy can fail in patients with biologically aggressive disease that do not benefit from resection. Therefore, an accurate molecular characterization of tumor phenotype will help in predicting prognosis and chemotherapy sensitivity, as well as inform decisions regarding upfront resection and the most appropriate drug choice for chemotherapy. Deep tumor molecular profiling constitutes an important source of information regarding tumor phenotype and biology, with impact on the choice of available therapeutic strategies. This information will increase the likelihood of success and also spare patients from unnecessarily aggressive therapeutic interventions. The goal of this study was to identify a molecular signature based on the transcriptomic profiles of PDAC patients that would allow for prediction of tumor progression and response to therapy.

To obtain an unbiased predictor of tumor aggressiveness, we established a series of patient-derived xenografts (PDX) from a multi-centric clinical trial that included resectable, locally advanced and metastatic PDAC patients. From these PDX samples, a transcriptomic signature (indicated as pancreatic adenocarcinoma molecular gradient; PAMG) was developed that accurately predicted tumor aggressiveness and resistance to mFOLFIRINOX, and could be applied to small amount of fine needle biopsies from EUS and formalin-fixed paraffin-embedded obtained tissue.

## Materials and Methods PaCaOmics patient’s cohort

Seventy-six patients with a confirmed PDAC diagnosis were included in this study. Clinical data was collected until July, 2017 (supplementary Tables I and II). Tumor samples were obtained from pancreatectomy in 40 patients (52.6%), EUS-FNA in 25 patients (32.9%) and carcinomatosis or liver metastasis during explorative laparotomy in 11 patients (14.5%). All samples were xenografted in immunocompromised mice producing PDX samples.

### BACAP patient’s cohort

The BACAP (Base Clinico-Biologique de l’Adénocarcinome Pancréatique) cohort is a prospective multicenter pancreatic cancer cohort (ClinicalTrials.gov Identifier: NCT02818829. Registration date: June 30, 2016) with a biological clinical database. Treatment naive tumor biological samples from endoscopic ultrasound-guided fine-needle aspiration (EUS-FNA) were available for 60 patients. Survival analysis was performed on the 47 patients with locally-advanced or metastatic diseases that subsequently received chemotherapy.

### Transcriptomic profiling and analysis

RNA was obtained from all PDX and BACAP cohort samples, for more details see supplementary matherial and methods. Next Generation Sequencing (RNA-seq) was performed on these samples and assessed. Details on transcriptomic profiling and analysis are available in the supplementary information. The PAMG is available as an online application (http://cit-apps.ligue-cancer.net//pancreatic_cancer/pdac.molgrade) and as an R package (https://github.com/RemyNicolle/pdacmolgrad).

## Results

### Using PDX to define the molecular diversity of PDAC

Recent reports indicate PDAC can be classified into distinct, biologically relevant categories based on histological and molecular analysis [4, 5]. However, relatively few patients (15%) undergo resection that allows this analysis, and high intra-tumor heterogeneity and the limited amount of material obtained from EUS-FNA diagnostic biopsies prevent a precise classification of all PDAC tumors. One solution to circumvent these problems is transplantation of PDAC tumors into immunodeficient mice to produce patient-derived xenografts (PDX). This process makes it possible to obtain PDXs from EUS-FNA diagnostic biopsies providing adequate material to determine PDAC histological classes for locally advanced or metastatic tumors. We observed that PDXs are less complex and heterogeneous tumors, but faithfully recapitulate the molecular profiles and histology of the original patient tumors. Another important point that conduct us to choose PDX as model is that it offers the posibility to distinguish between the tumor and stromal cells. In fact, sequencing profiles of a mix of human grafted cancerous and infiltrating mouse stromal cells can be analyzed separately *in silico* by unambiguously assigning each sequence to the human or mouse genome [6]. Therefore, we generated PDX samples for a cohort of patients (PaCaOmics) to define histological and molecular grades for each sample.

First, we assessed the histology of PDX using the entire cohort of 76 patients. PDX were ranked into five different histological classes by two blinded expert pathologists ranging from the less differentiated PDX (class I), which is associated with the most aggressive phenotype, to the most differentiated PDX (class V; Figure S1). The here described five histological classes of PDX strongly correlates with the expression of genes defining the already described molecular subtypes [6-9] as higher expression of genes linked to the classical PDAC subtype is correlated with increased differentiation of PDX samples, combined with lower expression of genes linked to basal-like subtype (Figure 1a and Figure S1). Interestingly, the variation in the expression of the classical genes towards the basal-like genes vary gradually from the more differentiate to the less differentiate histological classes respectively. Therefore the precise histological analysis of PDX suggests that molecular classification of PDAC is more complex than a two-class dichotomy (i.e.basal-like and classical). We next employed a consensus clustering approach on whole-transcriptome with increased subtypes splitting. Figure 1b shows the clustering results in 2 to 4 subtypes which, similarly to histological classification, demonstrate a gradual increase and decrease in genes of the classical and basal-like subtypes respectively.

**Figure 1.**
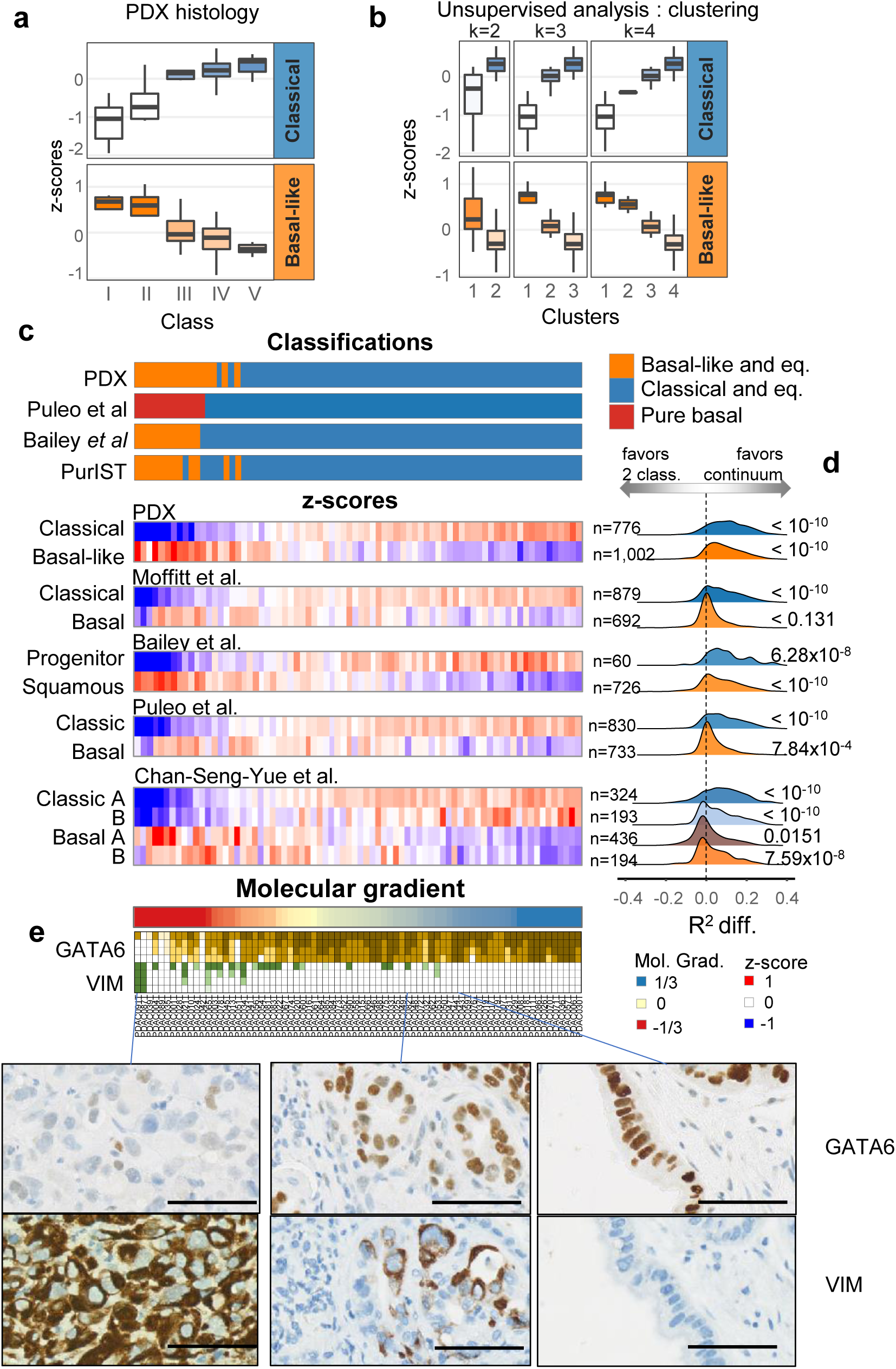
PDAC gene signatures and classification in PDX. **a.** Normalized and averaged expression of genes specific to the classical and basal-like subtypes in PDX (n=76) grouped by a five-subtype histological classification. **b.** Unsupervised classifications in k classes by consensus clustering (with k from 2 to 4) and association of each cluster to basal-like and classical gene expression. On a. and b. boxplots are colored by the median z-score of each group. **c.** Heatmap representation of the transcriptomic characterization of the PDX (n=76) with each PDX as a column. Previously published classifications were applied to the human transcriptome profiles of the PDX. Non-tumor driven classifications were applied (ADEX, Immunogenic, desmoplastic, activated stroma, Immune classical), however, no PDX were assigned to any of them. The z-score of each of the published classification gene sets is represented. The number of genes of each signature is annotated on the right of the heatmap. PDX were ordered by their value on the molecular gradient. **d.** Distribution of the differences in the coefficient of determination (R2) between two generalized linear models associating the expression of each gene in each signature with either the two-class classification from PurIST or the Molecular Gradient. The distribution of R2 differences was compared to that of other genes (not found in any other subtype signatures) using Welch’s t test. **e.** GATA6 and Vimentin (VIM) immunohistochemical quantification. Four levels of staining were used to quantify the proportion of cells at each four levels of GATA6 or VIM protein expression.

Histological and molecular classifications of PDX suggest PDAC diversity may be better represented by a continuum of differentiation that is as also followed at the molecular level. To establish a robust continuous molecular description of PDAC, we applied an unsupervised approach termed independent component analysis (ICA) previously shown to derive highly reproducible signatures from transcriptome profiles by extracting biologically relevant components [10, 11]. Figure S2 illustrates the procedure used to uncover an RNA signature which, in essence, builds on the blind deconvolution of the PDX transcriptomic profiles to generate component spaces. The component (and its associated space) that best correlated to the PDX histological classification was selected and, in analogy to histological grading, was termed the pancreatic adenocarcinoma molecular gradient (PAMG). The PAMG is computed from a weighted combination of gene expression values, standardized around zero with non-outlier values between -1 and +1. Figure 1c shows the different molecular classifications of PDAC applied to the 76 PDX along with the summarized expression of each of the previously proposed subtypes. To evaluate whether a continuous or dichotomous description of PDAC epithelial diversity is more relevant, gene expression in each of these signatures was fitted with the proposed PAMG and with the latest basal-like/classical classifier PurIST. The difference in the coefficient of determination (R2) of the two models was compared to the background (genes not in any of the assessed signatures, n=7,393) showing overall that a continuum is likely to be a more reliable description of PDAC molecular diversity. We observed that PAMG produces a better description on 11 out of 12 signatures tested by a Welch’s t test (Figure 1d).

A continuum of phenotypes would predict that extreme cases would be more homogeneous, composed of highly specified epithelial cells from the corresponding end of the spectra (*i.e.* basal-like or classical). PDAC cases in the middle of the spectrum could either be the result of a homogeneous intermediate epithelial phenotype or a mixture of extreme phenotypes of which bulk tumor analysis would result in an intermediary phenotype. To evaluate these non-mutually exclusive hypotheses, we performed immunostaining for GATA6, which we previously showed to be a major driver of the classical phenotype [12], and vimentin (VIM) in a tissue microarray containing all 76 xenograft tumors. VIM is a marker of mesenchymal differentiation and carcinomas with more aggressive behavior and poor histological differentiation [13]. Figure 1e shows quantitative results and representative examples of expression. While some GATA6+/VIM+ stained tumors exist, we generally observed a continuum of differentiation defined by increases in the level and proportion of expression of GATA6 along the PAMG that correlated with increased differentiation. Conversely, we observed VIM expression increasing gradually towards low differentiated phenotypes.

### Reproducibility of the PAMG in resectable human primary PDAC

To evaluate the robustness of the PAMG, we tested whether an equivalent RNA signature could be blindly reproduced in independent PDAC series with transcriptomic data. Two large series of PDAC were used for this purpose 269 resected tumors from the Australian ICGC [14] profiled on Illumina microarrays from frozen samples, and the multi-centric cohort of 309 consecutive patients from Puleo *et al.* [7], profiled on Affymetrix arrays from paraffin-embedded samples. To assess the reproducibility of the PAMG in these series of samples, a blind deconvolution of the transcriptomes was performed using ICA with increasing number of components resulting in ICA spaces of up to 25 unsupervised independent components (Figure 2a). Once components were extracted, a component matching the PAMG from the PDX was sought by correlating gene weights of both the reference PDX ICA space and the new ICA spaces to be evaluated. This analysis aimed at evaluating whether a component biologically similar to the PAMG could be extracted from the human tumor datasets. A molecular component equivalent to the PDX-derived PAMG was found in virtually all ICA component spaces in both datasets despite the difference in measurement technologies and in tissue preservation (Figure S3). The component with the highest gene weight correlation to the PDX-based PAMG was selected from each dataset. Figure 2b illustrates the overall consistency in the gene weights defining each of the components of three PAMGs. Overall, three components were selected from an unsupervised gene-expression deconvolution analysis applied to three independent datasets representing diverse technological (microarrays and RNAseq) and tissue (FFPE, Frozen, PDX) options to profile PDAC resulting in three biologically equivalent implementations of the PAMG.

**Figure 2.**
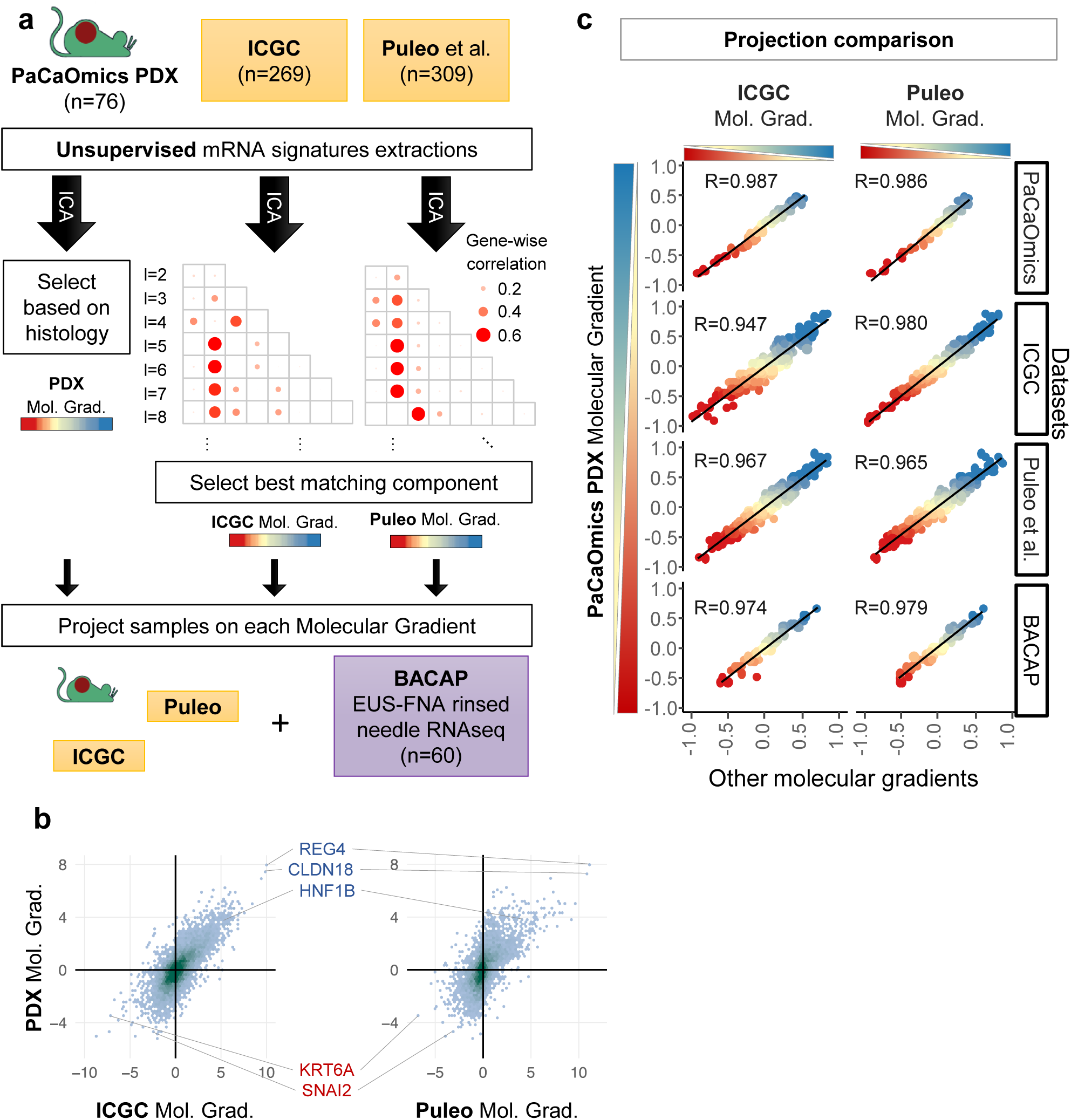
Reproducibility of the PAMG in PDAC. **a.** Schematic illustration of the identification of the PAMG in public datasets. ICA (independent component analysis) blind deconvolution was used on three different datasets of whole transcriptome profiling, generating spaces of independent components of increasing sizes (2 ≤ *l* ≤ 25). The PAMG was first obtained from PDX by selecting the component most associated with PDX histology. The gene weights of this initial PDX-based independent component was then correlated to the gene weights of all extracted independent components in the other datasets, with the spearman correlation represented in a grid. The highest correlating component of each dataset was selected as the PAMG. **b.** Density plot of the PAMG gene weights of common genes found in each pair of datasets. Marker genes are highlighted. **c.** Scatter plots comparing the three versions of the Molecular Gradient (PDX, ICGC and Puleo) on four datasets. Each point is a sample, colored by its PAMG score as defined by the PDX version. Pearson correlation is shown.

While the three independently identified PAMGs share a similar gene expression basis, we next sought to evaluate the extent they define the same PDAC heterogeneity. The samples from the three different datasets were each projected on all three PAMGs. Figure 2c shows a high correlation between the three PAMGs in all three datasets, demonstrating that the signatures measure a common biological diversity independent of the types of samples profiled and the technologies used. To validate this high reproducibility, the same analyses were applied to a PDAC cohort consisting of 60 RNAseq profiles from RNA obtained by rinsing EUS-FNA diagnostic biopsies. The three versions of the PAMGs gave highly similar results on FNA-derived samples (R>0.97).

### The PAMG is associated to tumor aggressiveness

Several studies using only resectable tumors show molecular diversity of the epithelial compartment of PDAC is associated with tumor aggressiveness and patient prognosis [7, 8, 14]. Our next goal was to determine if the PAMG could be predictive in all PDAC tumors. To assess the prognostic value of the PAMG, association with overall survival was first evaluated on the ICGC series [14] which consisted of 267 resected patients with follow-up, and 230 samples with histological characterization. The continuous value of the PAMG (as extracted from the ICGC transcriptome dataset) was strongly associated to patient’s overall survival (univariate Hazard Ratio: uHR=0.405, [0.255-0.642]; Wald P-value: p=1.23×10^−4^) and compared favorably to the basal-like/classical dichotomous classification (Figure 3a and Figure S4). A virtually identical result was obtained with the other PAMGs derived from the PDX and Puleo *et al.* cohorts (Figure S4). The continuous characterization of patients in the ICGC series by the PAMG showed a positive correlation with significant increase in OS (Figure 3b) also illustrated in a Kaplan-Meier analysis (Figure 3c) after splitting the PAMG using three arbitrary thresholds (−0.5, 0 and 0.4; selected on the basis of the separation of histological classes of PDX). Importantly, a weak association was found between the PAMG and the histological differentiation of these tumors (Figure S4), suggesting a partial relationship between molecular classification of PDAC and traditional histological classes [7]. In a multivariate analysis including the PAMG and the histology of these tumors, the PAMG was an independent predictor of OS (Figure 3d).

**Figure 3.**
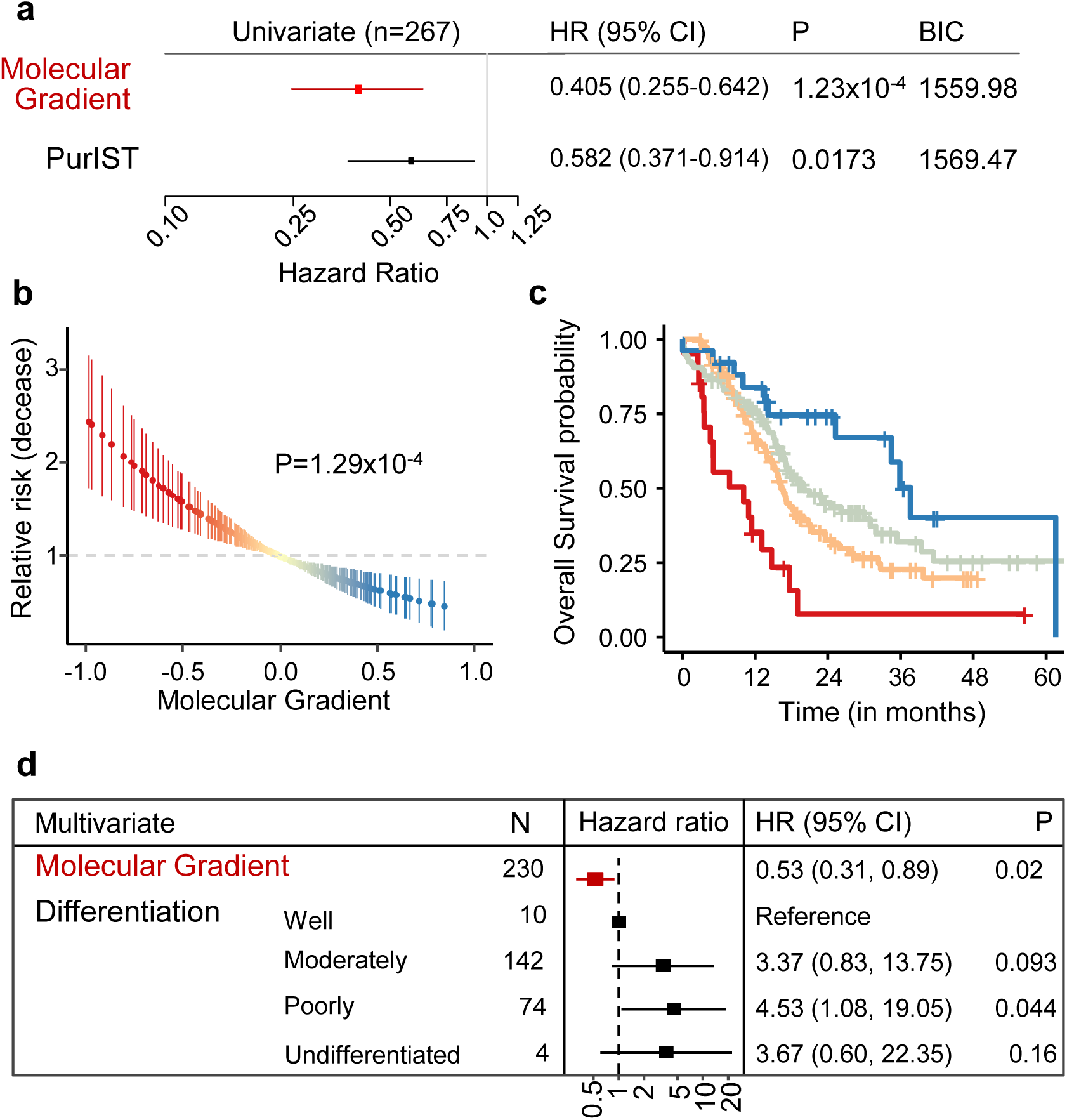
Prognostic value of the PAMG in the ICGC series. **a.** Univariate survival analysis using the overall survival (OS) of 260 patients associated with either the PAMG or the PurIST two-subtype classification. **b.** Univariate relative risk for OS associated with the PAMG. Each point is a patient’s relative risk of disease with error bars corresponding to a 95% confidence interval. **c.** Kaplan-Meier plot of survival using arbitrary cuts of the Molecular Gradient. d. Multivariate survival analysis forest plot. Univariate: n=267. Multivariate: n=230. Wald’s test p-values are shown.

To further assess the value of the PAMG in a more reliable cohort of patients, the multi-centric cohort of 309 consecutive patients from Puleo *et al*. [7] was used. This very complete cohort contains whole follow up for 308/309 patients (median follow-up 51.4 months) and with a majority (298/309) also having data on extended clinical and pathological characterization. The PAMG was associated with patients OS (uHR = 0.321, [0.207-0.5] ;p=4.97×10^−7^) and compares favorably to the basal-like/classical classification (Figure 4a and Figure S5). The PAMG was correlated to a positive outcome in Puleo cohort, with a progressive improvement of OS coinciding with higher PAMG levels (Figures 4b and 4c). A multivariate analysis including resection margins, histological grading and TNM Node status demonstrated the PAMG is an independent prognostic factor in resected PDAC (Figure 4d).

**Figure 4.**
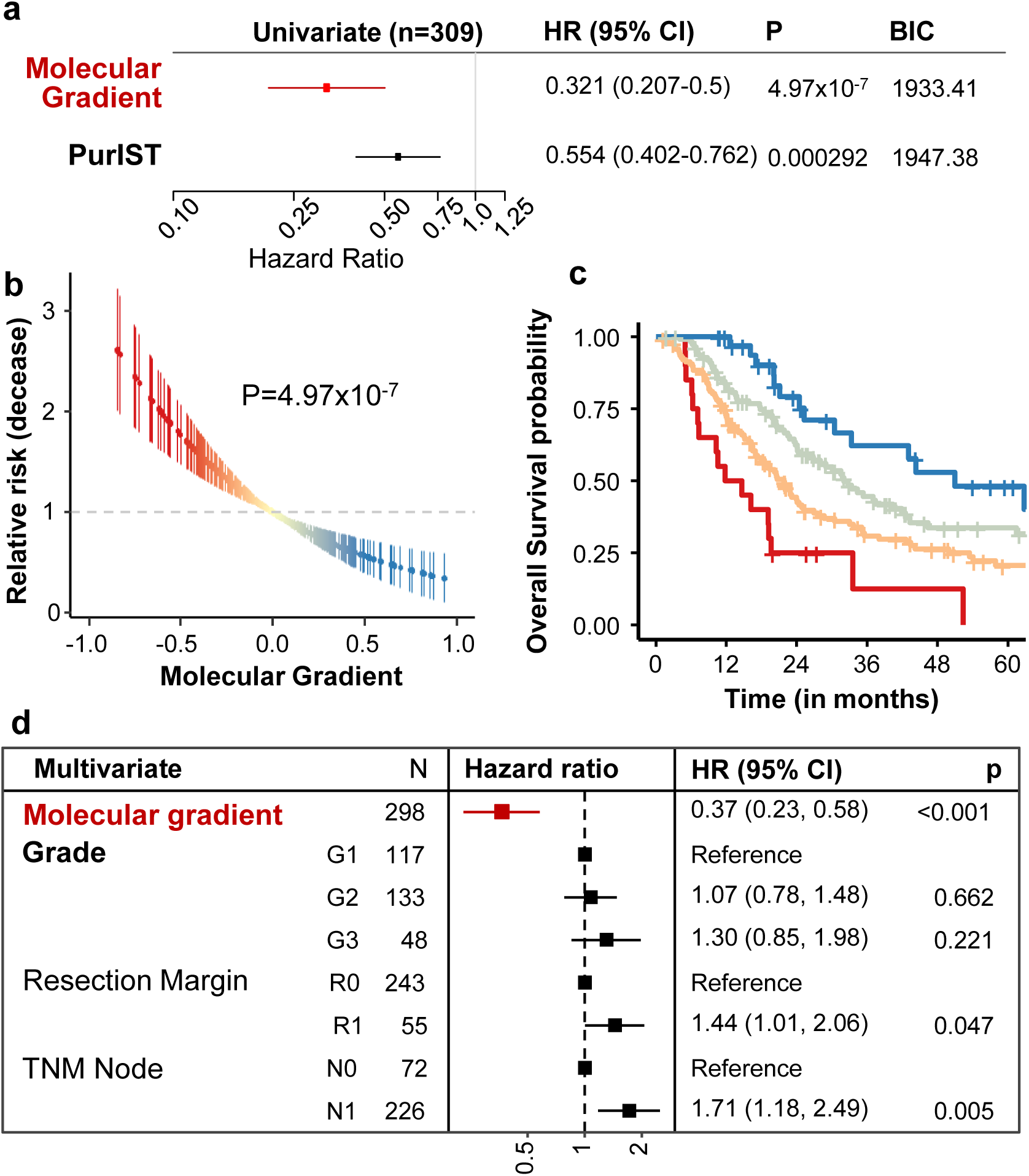
Prognostic value of the PAMG in the Puleo cohort. **a.** Univariate survival analysis using the OS of 308 patients associated with either the PAMG or the PurIST two-subtype classification. **b.** Univariate relative risk for OS associated with the PAMG. Each point is a patient’s relative risk of decease with error bars corresponding to a 95% confidence interval. **c.** Kaplan-Meier plot of survival using arbitrary cuts of the PAMG. **d.** Multivariate survival analysis forest plot. Univariate: n=308. Multivariate: n=298. Wald’s test p-values are shown.

### PAMG predicts the clinical outcome of advanced PDAC patients

The clinical relevance of the PAMG is dependent on its applicability to work on biopsy samples obtained prior to treatment. In the BACAP cohort, RNA was extracted from 60 samples obtained by rinsing the echoendoscopy-guided fine needles. The original aspirate was used for diagnosis. Figure 2c shows all three versions of the PAGM gave the same result on these small sample biopsies. The PAGM was also associated with the OS of the 47 patients with advanced diseases (uHR=0.308, [0.113-0.836]; p=0.0208, Figure 5a) and, similar to resectable tumors, compared favorably to the PurIST two-subtype classification. The PAGM was also associated to survival in a multivariate model including the tumor stage (Figure 5b).

**Figure 5.**
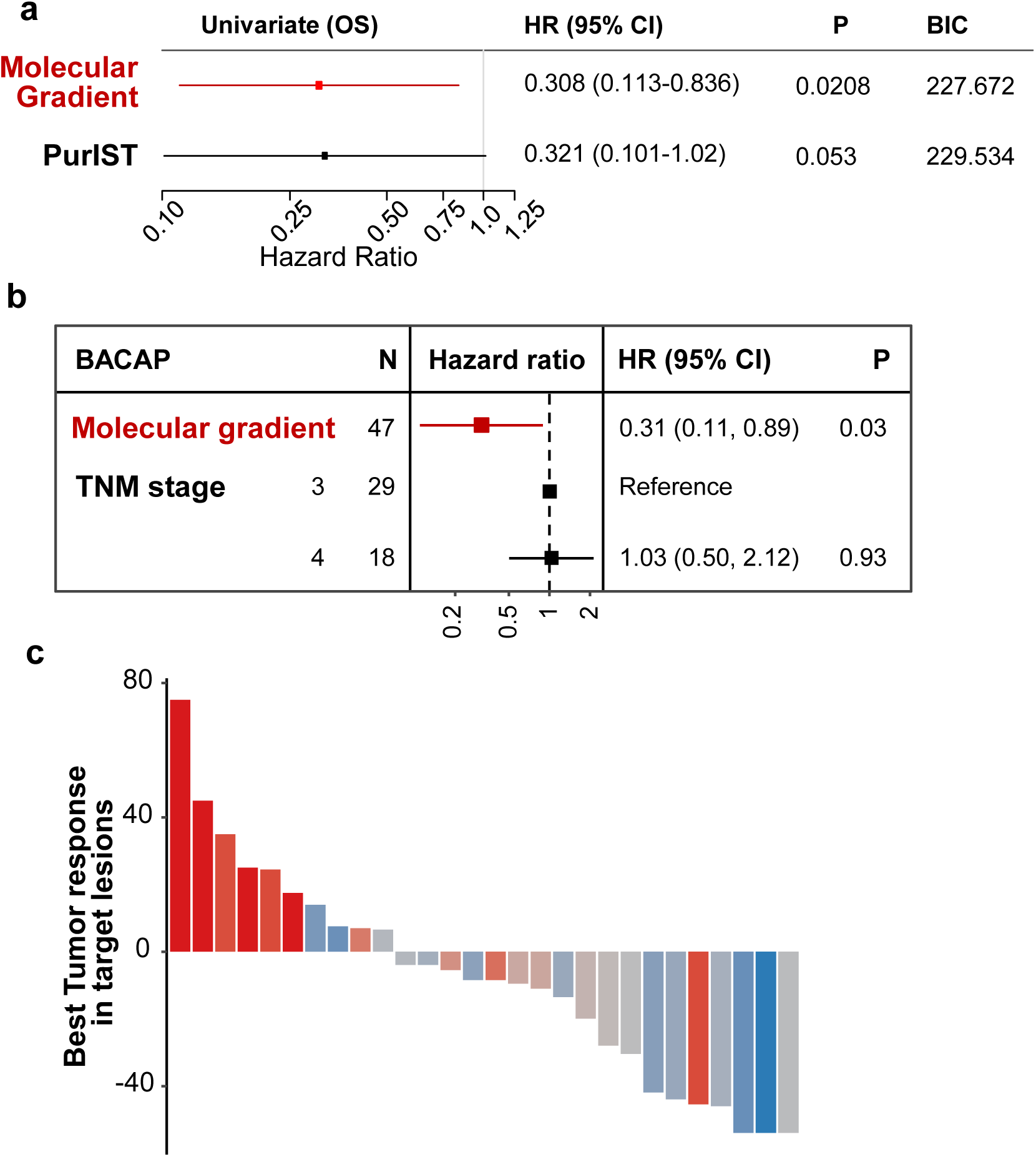
Evaluation of the PAMG in advanced disease. **a.** Univariate survival analysis using the OS of 47 patients in the BACAP cohort associated with either the PAMG or the PurIST two-subtype classification. **b.** Multivariate survival analysis forest plot for the BACAP cohort. **c.** Waterfall plot illustrating the change in tumor size induced by mFOLFIRINOX treatment evaluated by RECIST 1.1 in the COMPASS cohort (n=28). Annotated Pearson’s correlation between RECIST 1.1 and PAMG is shown.

### PAMG predicts the response to mFOLFIRINOX of advanced PDAC patients

It was previously suggested that molecular subtypes of PDAC were associated with responses to chemotherapy, in particular FOLFIRINOX [8, 15, 16]. Therefore, to evaluate the predictive value of the PAGM to chemotherapy response, it was applied to metastatic patients in the COMPASS trials for which transcriptomic profiles and tumor responses to mFOLFIRINOX were available [16]. The objective response was significantly associated with the PAMG (Figure 5c, R= -0.67; p < 0.001), with more aggressive tumors (i.e. low on the PAMG) showing little to no response to mFOLFIRINOX.

## Discussion

An important factor in determining treatment options for PDAC involves the ability to accurately classify the tumor and predict the aggressiveness of the disease. However, resolving the diversity of molecular tumor phenotypes in PDAC is a complex issue involving the necessary distinction of transformed and non-transformed cells as well as a multi-scale integration in which microscopic cellular phenotypes are considered with macroscopic phenotypes of the whole-tumor tissue. Previous work has mainly focused on resected primary PDAC tumors, often resulting in classifications that considers all of the cell types within the tumor (*e.g.* the infiltrated Immunogenic subtype), and delineated a consensual basal-like versus classical dichotomy. However, this two-subtype classification system of PDAC has recently been challenged by several studies showing the coexistence of basal-like and classical cells in the same tumors as well as to the likely existence of intermediate cellular phenotypes [15, 17]. In this study, we have used a gradient system that takes this into consideration to classify PDAC. The resulting PAMG signature is more informative and clinically relevant than a binary non overlapping method.

Single cell RNA sequencing [15] and immunohistochemistry [18] of PDAC revealed intra-tumor heterogeneity where both types of cells (basal-like and classical) frequently co-exist. RNA profiling of multiple regions or multiple lesions from the same patient also demonstrated intra-tumor heterogeneity of the transformed epithelial compartment [4]. Using single cell RNA-seq, Chan-Seng-Yue *et al*. in 2020 [15] confirmed the presence of several subpopulations with differential proliferative and migratory potentials in PDAC. In particular, they observed two ductal subtypes with abnormal and malignant gene expression [15]. Our own unpublished results identified four common cell clusters in patients with a classical PDAC. These four clusters were present in different proportions in all tumors examined, with one of these clusters corresponding to a basal-like phenotype, even though the tumors were classified as classical by global RNAseq analysis.

We have made similar observations in this study. VIM, which is mainly expressed in basal-like subtype, was detected by immunohistochemistry in almost all classical tumors, with variable levels of expression [17]. We detected few VIM+ cells in tumors presenting an intermediate PAMG. In other words, very classical or very basal-like subtypes are mainly composed by pure cells, but the intermediate subtype is the consequence of a mix of classical and basal-like subtypes and/or an intermediate phenotype. These observations question the relevance of a dichotomous model of PDAC diversity and makes the molecular description a different and complex scenario for every tumor. Since PDAC tumors are heterogeneous, this must be taken into consideration for classification and treatment purposes Protocols characterizing the proportion of intermediate cell types or tumor heterogeneity are necessary.

In this work, we developed a molecular gradient that defines a continuum of PDAC phenotypes. We developed 76 PDX, obtained from resectable and unresectable PDAC, since they offer a platform with an incomparable discrimination of transformed and non-transformed cells RNA. First, we applied a deconvolution algorithm (ICA) to the transformed epithelial RNA profiles to identify in an unsupervised manner the RNA signatures that best defined the heterogeneity of PDX and, in particular, its aggressiveness. This approach extracted a specific RNA signature robustly identified in PDX and human primary tumors with a minor effect of tissue preservation (FFPE vs. frozen), RNA profiling platform (microarrays or RNAseq) or of the algorithm’s parameter (the total number of extracted components). This RNA signature, termed PAMG, provides a score measuring the molecular level of differentiation of a given sample derived from a whole-transcriptome profile. The approach for phenotyping is robust since several signatures extracted from different datasets gave highly similar results.

The PAMG introduces a simple framework, based on a simple RNA signature compatible with all previously proposed PDAC classifications. The genes previously described as defining PDAC subtypes were in fact better explained by the PAMG than by the two-class classification itself. Molecular classifications of PDAC and, in particular, the basal-like/classical dichotomy, is a major prognostic factor in most datasets and is typically shown to correlate with response to FOLFIRINOX. Our results showed the PAMG holds superior clinical value that could be ascertained prior to entering any curative protocols, using any current diagnostic material including EUS-guided biopsy needle flushing. This model could have a major impact on patients who are cleared for resection by identifying patients that will have an unfavorable disease evolution and may benefit from initial neoadjuvant therapy prior to upfront surgery. Another kind of patient the PAMG could impact are the 20 to 30% percent of patients diagnosed with a locally-advanced disease. If pancreatectomy and simultaneous arterial resection has traditionally been considered as a general contraindication to resection [19], some of these patients with good prognosis might indeed benefit from aggressive surgical approaches [20].

In conclusion we propose a transcriptomic signature that unifies all previous molecular classifications of PDAC under a continuous gradient of tumor aggressiveness that can be performed on FFPE samples and EUS-guided biopsies. In addition to its strong prognostic value, it may predict mFOLFIRINOX responsiveness.

## Acknowledgments

This work is part of the national program Cartes d’Identité des Tumeurs (CIT) funded and developed by the Ligue Nationale Contre le Cancer. This work was supported by INCa (Grants number 2018-078 and 2018-079, BACAP BCB INCa_6294), Canceropole PACA, DGOS (labellisation SIRIC), Amidex Foundation, Fondation de France and INSERM. Authors wish to thanks Christopher Pin for critically reading the manuscript and IPC/CRCM Experimental Pathology platform for TMA and immunohistochemistry performing. This research was supported by the CRCM Integrative Platform (Cibi) and the CRCM’s DataCentre for IT and Scientific Computing (DISC).

## The BACAP consortium

1 Barbara Bournet, Cindy Canivet, Louis Buscail, Nicolas Carrère, Fabrice Muscari, Bertrand Suc, Rosine Guimbaud, Corinne Couteau, Marion Deslandres, Pascale Rivera, Anne-Pascale Laurenty, Nadim Fares, Karl Barange, Janick Selves, Anne Gomez-Brouchet. 2 Bertrand Napoléon, Bertrand Pujol, Fabien Fumex, Jérôme Desrame, Christine Lefort, Vincent Lepilliez, Rodica Gincul, Pascal Artru, Léa Clavel, Anne-Isabelle Lemaistre. 3 Laurent Palazzo ; 4 Jérôme Cros ; 5 Sarah Tubiana ; 6 Nicolas Flori, Pierre Senesse, Pierre-Emmanuel Colombo, Emmanuelle Samail-Scalzi, Fabienne Portales, Sophie Gourgou, Claire Honfo Ga, Carine Plassot, Julien Fraisse, Frédéric Bibeau, Marc Ychou ; 7 Pierre Guibert, Christelle de la Fouchardière, Matthieu Sarabi, Patrice Peyrat, Séverine Tabone-Eglinger, Caroline Renard; 8 Guillaume Piessen, Stéphanie Truant, Alain Saudemont, Guillaume Millet, Florence Renaud, Emmanuelle Leteurtre, Patrick Gele ; 9 Eric Assenat, Jean-Michel Fabre, François-Régis Souche, Marie Dupuy, Anne-Marie Gorce-Dupuy, Jeanne Ramos ; 10 Jean-François Seitz, Jean Hardwigsen, Emmanuelle Norguet-Monnereau, Philippe Grandval, Muriel Duluc, Dominique Figarella-Branger ; 11 Véronique Vendrely, Clément Subtil, Eric Terrebonne, Jean-Frédéric Blanc, Etienne Buscail, Jean-Philippe Merlio ; 12 Dominique Farges-Bancel, Jean-Marc Gornet, Daniela Geromin ; 13 Geoffroy Vanbiervliet, Anne-Claire Frin, Delphine Ouvrier, Marie-Christine Saint-Paul; 14 Philippe Berthelémy, Chelbabi Fouad ; 15 Stéphane Garcia, Nathalie Lesavre, Mohamed Gasmi, Marc Barthet ; 16 Vanessa Cottet ; 17 Cyrille Delpierre.

1 The CHU and the University of Toulouse, Toulouse, France; 2 Jean Mermoz Hospital, Lyon, France; 3 Trocadéro Clinic, Paris, France; 4 The Department of Pathology, Beaujon Hospital and Paris 7 University, Clichy, France.5 The Biobank, Bichat Hospital and Paris 7 University, Paris, France; 6 The Cancer Institute and the University of Montpellier, Montpellier, France; 7 The Léon Bérard Cancer Center, Lyon, France;8 The Department of Digestive Surgery, the CHU and the University of Lille, Lille, France;9 The CHU and the University of Montpellier, Montpellier, France.10 La Timone Hospital and the University of Marseille, Marseille, France;11 The CHU and the University of Bordeaux, Bordeaux, France;12 Saint Louis Hospital and Paris 7 Diderot University, Paris, France;13 The CHU and the University of Nice, Nice, France;14 Pau Hospital, Pau, France;15 The CHU Nord Hospital and the University of Marseille, Marseille, France;16 INSERM UMR866 and the University of Dijon, Dijon, France;17 INSERM UMR1027 and the University of Toulouse, Toulouse France.

## Author contributions

**Conception and design:** Nelson Dusetti, Rémy Nicolle and Juan Iovanna

**Provision of study materials or patients:** Marine Gilabert, Vincent Moutardier, Stephane Garcia, Pauline Duconseil, Charles Vanbrugghe, Philippe Grandval, Marc Giovannini, Cindy Canivet, Barbara Bournet and Louis Buscail.

**Histological classification and analysis:** Nicolas Brandone, Flora Poizat, Marion Rubis; Jérôme Cros.

**Collection and assembly of data:** Martin Bigonnet, Odile Gayet, Julie Roques, Pauline Duconseil, Nabila Elarouci, Lucile Armenoult, Mira Ayadi and Aurélien de Reyniès.

**Data analysis and interpretation:** Nelson Dusetti, Rémy Nicolle, Yuna Blum, Samir Dou and Juan Iovanna.

**Manuscript writing:** Nelson Dusetti, Rémy Nicolle, Yuna Blum and Juan Iovanna

**Final approval of manuscript:** All Authors

## Competing financial interest

The authors declare no competing financial interest

## Legend of Supplementary Figures

**Figure S1.**
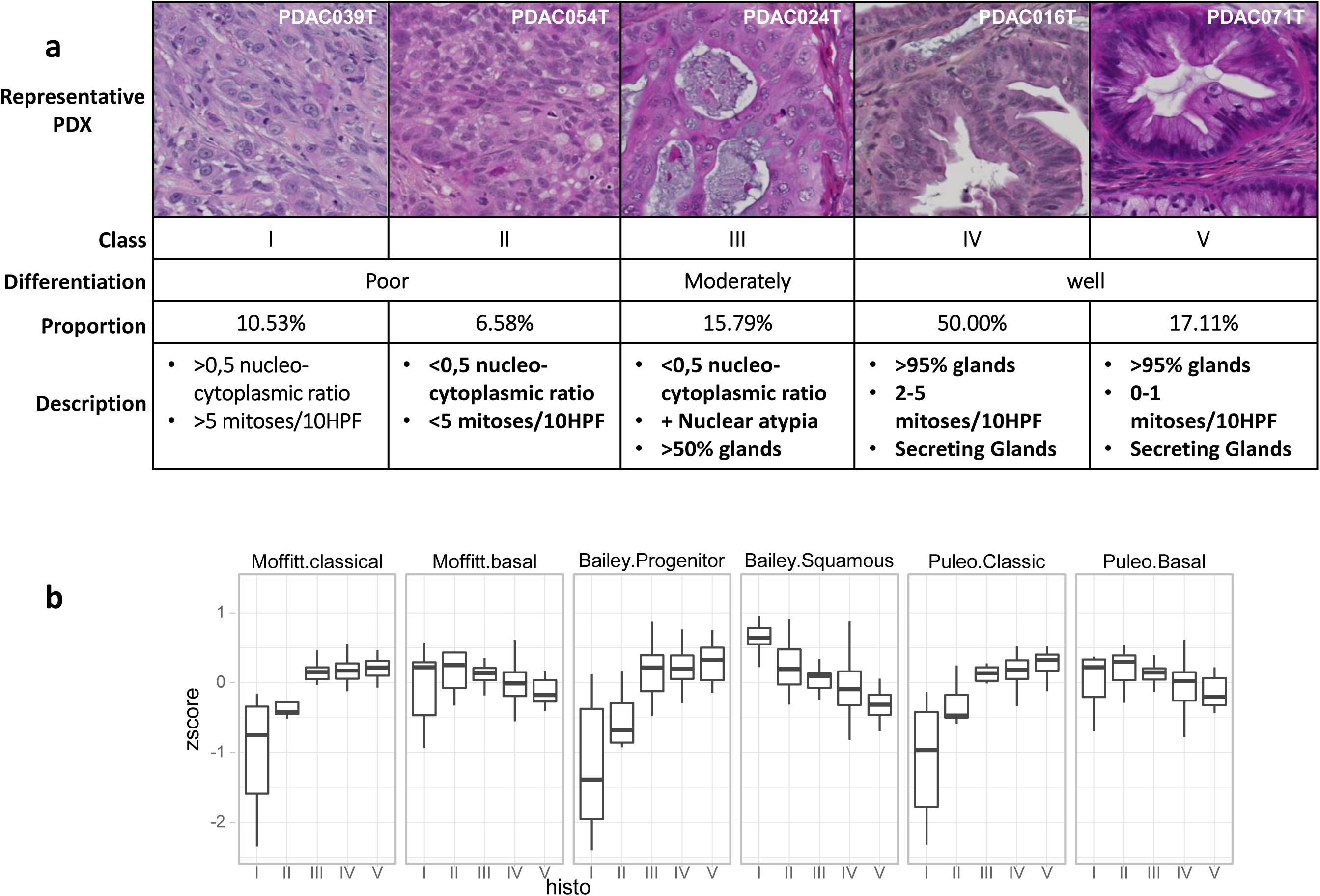
Histological classification of PDX. **a.** Representative examples for each histological class and the associated proportions within the poor-, moderate- and well-differentiated PDX in the PaCaOmics cohort (total n=76). HPF: High-power field. **b.** Distribution of the z-scores of each subtype-specific gene sets according to the five histological classes of PDX.

**Figure S2.**
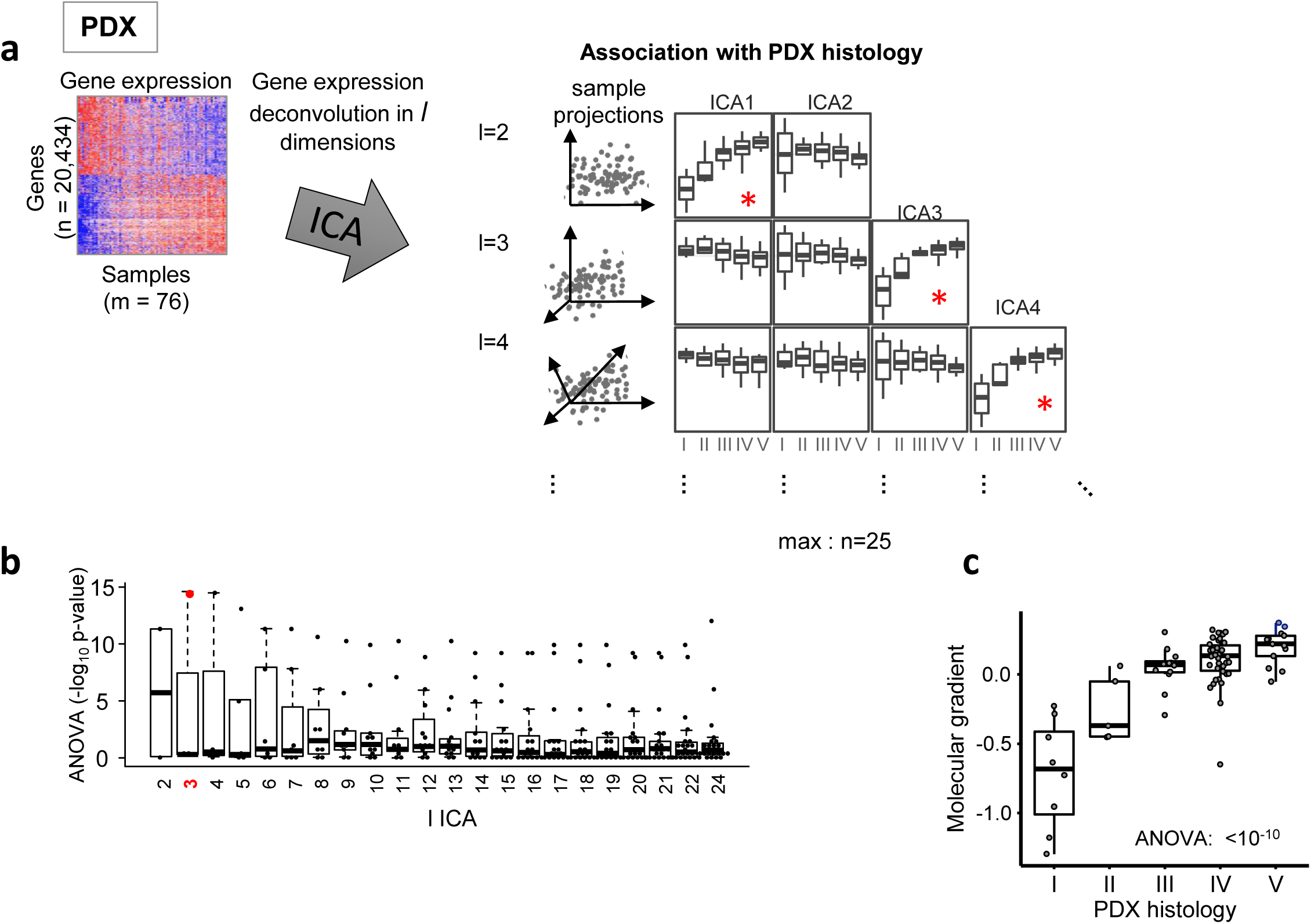
Identification of the Molecular Gradient from the transcriptomic profiles and the histological classes in PDX. **a.** Illustration of the process. ICA is applied to the 76 gene expression profiles generating ICA system with increasing number of components l. Each component of each ICA system are associated to the PDX histological classification. **b.** Distribution of the association between every component in each ICA system and the histological classification. –log10 transformation of the ANOVA p-value is represented. **c.** Distribution for the selected component to be used as the Molecular Gradient.

**Figure S3.**
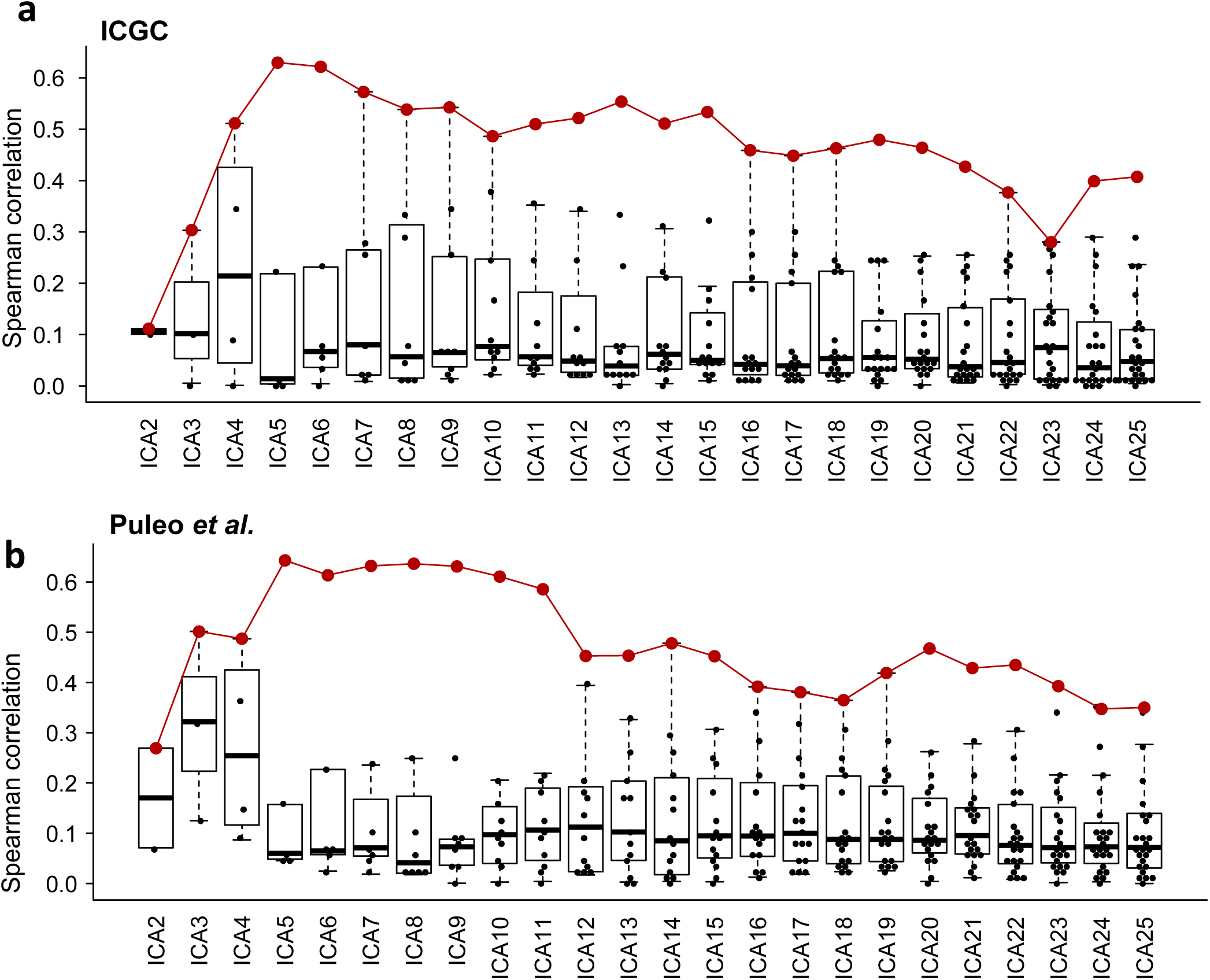
Reproducibility of the identification of the Molecular Gradient in human primary tumor series. Distribution of the Spearman correlation (absolute value) between the gene weights in the PDX-derived Molecular Gradient and every component of a given ICA system derived from the blind deconvolution of transcriptomic data from the ICGC series (**a**) and the Puleo et al. cohort (**b**). The maximum correlation, the best matching component, of each ICA system is shown in red.

**Figure S4.**
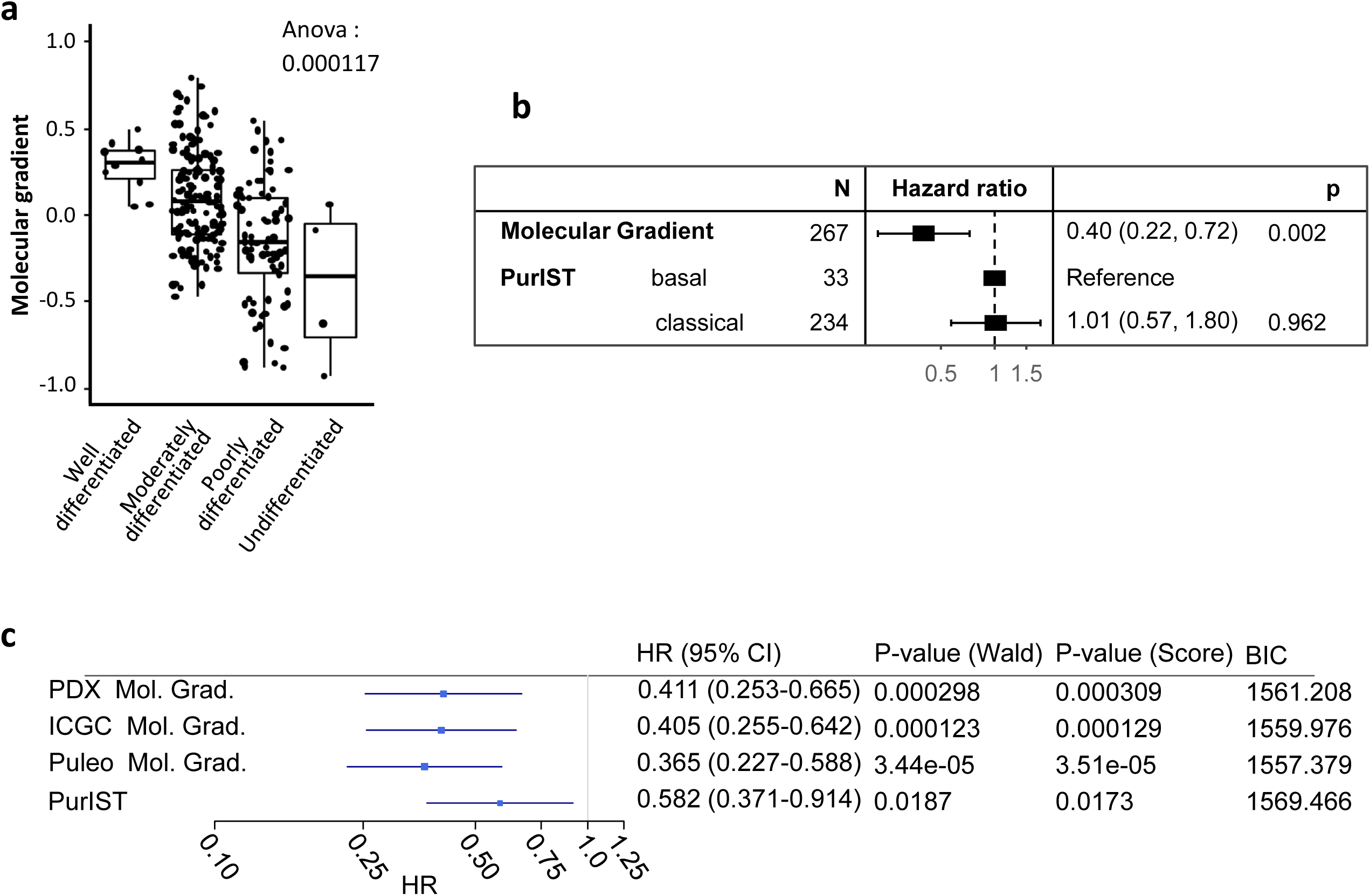
Additional analyses of the Molecular Gradient in the ICGC series. **a.** Association between Molecular Gradient and traditional histological characterization. **b.** Multivariate survival analysis including both the Molecular Gradient and the PurIST classification. **c.** Univariate survival analysis including all three versions of the Molecular Gradient.

**Figure S5.**
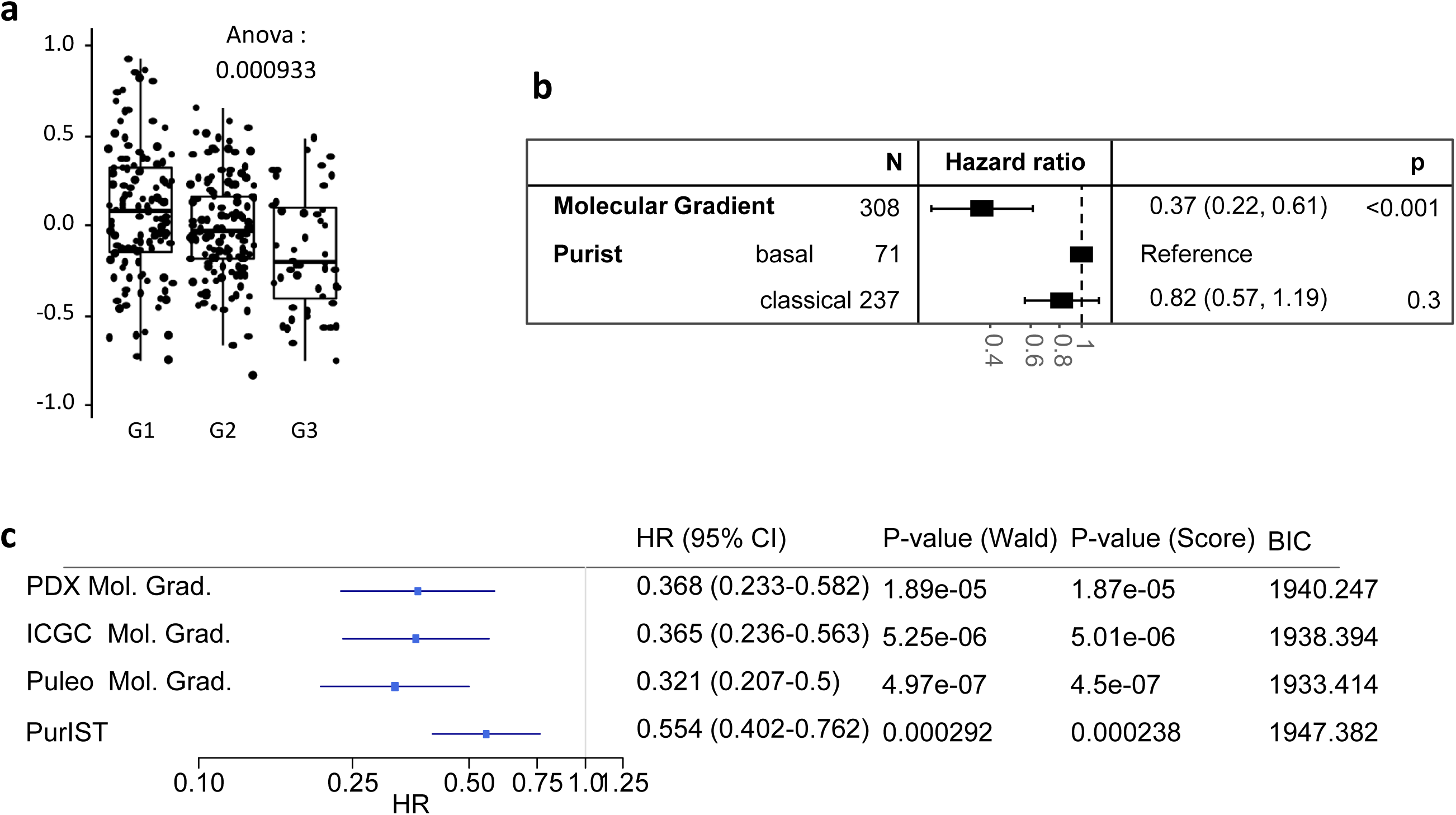
Additional analyses of the Molecular Gradient in the Puleo et al. cohort. **a.** Association between Molecular Gradient and traditional histological characterization. **b.** Multivariate survival analysis including both the Molecular Gradient and the PurIST classification. **c.** Univariate survival analysis including all three versions of the Molecular Gradient.

## Supplementary information

### PaCaOmics patient derived tumor xenograft and RNA-sequencing

All animal experiments were conducted in accordance with institutional guidelines and were approved by the “Plateforme de Stabulation et d’Expérimentation Animale” (PSEA, Scientific Park of Luminy, Marseille). Resected PDAC tissue was fragmented, mixed with 100 μL of Matrigel and implanted with a 10-gauge trocar (Innovative Research of America, Sarasota, FL) in the subcutaneous right upper flank of an anesthetized male NMRI-nude mouse (Swiss Nude Mouse Crl: NU(lco)-Foxn1nu; Charles River Laboratories, Wilmington, MA). Alternatively, samples obtained from direct tumor endoscopic ultrasound-guided fine needle aspiration (EUS-FNA) were mixed with 100 μL of Matrigel (BD Biosciences, Franklin Lakes, NJ) and injected as above. Once xenografts reached 1 cm3, they were removed and passed to NMRI-nude mice. After 3 passages, tumors were isolated and RNA extracted using the miRneasy mini kit (Qiagen). RNA-seq was performed as previously described [9, 20] using Illumina’s TrueSeq Stranded RNA LT protocol to obtain 100b paired-end reads. RNA-seq reads were mapped using STAR and SMAP on the human hg19 and mouse mmu38 genomes. Gene expression profiles were obtained using FeatureCount and normalized using the upper-quartile approach [21].

Tumor differentiation was defined based on the following established criteria, briefly: tumors were considered poorly differentiated when tissue architecture is solid, forming massive structures or with isolated cells without visible glandular structures in more than 50% of the tissue. This group included two classes (I and II) based on the degree of cyto-nuclear atypia and degree of mitosis. Class I tumors showed high nucleo-cytopasmic ratios (>0.5), and nuclei with irregular contours, dense chromatin, and/or prominent nucleolus. A high proportion of mitoses (>5 per 10 high-power field [HPF]) was also visible in this subgroup. Class II includes tumors with fewer atypia with a nucleo-cytoplasmic ratio < 0.5, regular-contoured nuclei, fine chromatin and a fine nucleolus. Mitoses were less frequent than in class I (< 5 mitosis/10 HPF). Class III includes tumors that were moderately differentiated with both types of architectures, glands made up 50-95% of the tumor, massive structures and nucleo-cytoplasmic atypia were less frequent (approximately 50% of nuclei) than in class I and II. Class IV and V were included in well differentiated PDX. They present a glandular architecture without solid component in more than 95%. In this group, class IV presents glands with cubic or short cylindrical cells with low or absent mucus secretion. The nuclei remain predominantly polarized and the atypia are more marked than in class V (looser chromatin, increase in the size of the nuclei when compared with class V). Mitoses were more frequent than in class V (2-5 mitosis / 10 HPF). Class V corresponds to the most differentiated tumors, the glands secrete mucin and cells present a cylindrical form, the nucleus was localized at the basal pole of the cell (polarized). Nuclei were small, with regular contours and mature chromatin without visible nucleolus. Mitoses were less frequent (0-1 mitosis / 10 HPF) that in class IV.

### PDX and TMA immunohistochemistry (IHC)

76 blocks of pancreatic ductal adenocarcinoma (PDAC) xenografts embedded in paraffin were selected to be included in the PDX-TMA. Each block was cut with a HM340E microtome with Niagara system. Hematoxylin eosin staining (HES) was performed on 3 µm thick sections to localize the tumor. A Minicore Tissue Arrayer was used to punch cores from the selected paraffin blocks, and distribute them in new blocks. Two cores of 0.6 mm diameter were used for each PDX. The PDX-TMA paraffin blocks were cut and stained (HES) to validate the tumor morphology of each core. Blocks were then stored at 4°C.

The GATA6 (AF 1700, R&D Systems) and Vimentin (V6389, Sigma) immunostaining were performed on 3µm thick serial sections of PDAC PDX tumors or for the PDAC PDX-TMA. The immunohistochemistry was carried out on the Ventana Discovery XT (IPC-CRCM Experimental Pathology Platform - ICEP, CRCM, Marseille). After deparaffinization, antigens retrieval was performed with Citrate-based buffer pH 6.0 (Cell Conditioner #2) for GATA6 and with tris-based buffer with a slightly basic pH (pH 8.0, Cell Conditioner #1) for Vimentin IHC. Primary antibodies were both incubated for 1 hour at 37°C. Then, an OmniMap anti-Goat HRP (HRP multimer) was used with DAB for GATA6 staining while a rabbit secondary antibody (Santa Cruz sc-454 at 1:500 dilution) was used before the appropriated OmniMap-HRP (anti-rabbit) for vimentin IHC. Finally, the counterstaining was done with hematoxylin and slides were cleaning, deshydrated and coversliped with permanent mounting media. GATA6 antibody was used at 1:40 dilution and Vimentin at 1:1200. GATA6 quantification was done in a score from 1 to 4 considering diferent intesities of positive staining and four different percetages of stained cells (1=1-24, 2=26-50, 3=51-75 and 4=76-100%).

### BACAP patient cohort and transcriptome profiling of EUS-FNA needle flushing and RNA-sequencing

The diagnostic EUS-FNA biopsy needle flushing of 60 advanced patients from the prospective BACAP cohort were obtained from the biorepository (the BACAP database is managed by the Montpellier Cancer Institute Data Center with the Clinsight® software). RNA was extracted using the Qiagen Allprep purification Kit® (Qiagen, Courtaboeuf, France). RNA-Seq libraries are performed with NEBNext® Ultra™ II Directional RNA Library Prep Kit for Illumina according to supplier recommendations (NEB). The capture is then performed on cDNA libraries with the Twist Human Core Exome Enrichment System according to supplier recommendations (Twist Bioscience).

After each EUS-FNA process, needle flushing is done as previously described [22, 23] in an Eppendorf cryovial containing 500 μl of RNAprotect Cell Reagent (Qiagen, Courtaboeuf, France) for subsequent DNA and RNA extraction using the Qiagen Allprep purification Kit (Qiagen, Courtaboeuf, France).

RNA-Seq libraries are performed with NEBNext Ultra II Directional RNA Library Prep Kit for Illumina according to supplier recommendations (NEB). The capture is then performed on cDNA libraries with the Twist Human Core Exome Enrichment System according to supplier recommendations (Twist Bioscience). First of all, an RNA quality control is performed on Fragment Analyzer (AATI) with the RNA kit (DNF-489) to check the integrity of the RNA profile. The protocol permits to convert total RNA into a library of template molecules of known strand origin. Then a capture of the coding regions of the transcriptome is performed and the resulting library is suitable for subsequent cluster generation and sequencing. Briefly, the RNA is fragmented into small pieces using divalent cations under elevated temperature. cDNA is generated from the cleaved RNA fragments using random priming during first and second strand synthesis and sequencing adapters are ligated to the resulting double-stranded cDNA fragments and enriched by 7 PCR cycles. The coding regions of the transcriptome are then captured from this library using sequence-specific probes to create the final library. For that purpose, 500 ng of purified Libraries are hybridized to the Twist oligo probe capture library for 16h in a singleplex reaction. After hybridization, washing, and elution, the eluted fraction is PCR-amplified with 8 cycles, purified and quantified by QPCR to obtain sufficient DNA template for downstream applications. Each eluted-enriched DNA sample is then sequenced on an Illumina HiSeq4000 as paired-end 75b reads. Image analysis and base calling is performed using Illumina Real Time Analysis (2.7.3) with default parameters. RNA-seq reads were mapped using STAR [24] on the human hg19 genome. Gene expression profiles were obtained using FeatureCount [25] and normalized using the upper-quartile approach.

### Bioinformatics and Statistical analysis

This section described all the analysis performed on the processed and normalized datasets.

### Classification and z-score of previously published signatures

A Gene expression classifier and subtype specific gene sets were identified for each of the previously published PDAC classification systems. The z-score of each gene set is defined as the average expression in a single sample of all the genes of a given gene set, after gene-wise zero-centering and unit variance scaling.

**PDX**: basal-like and classical gene sets were obtained from a differential analysis of the human (i.e. transformed epithelial cell compartment) RNAseq comparing the multiomics-based classification (supplementary table of the original article). A centroid based classifier with Pearson distance was used as described in the bladder and colorectal consensus classification studies [26, 27]. The number of genes in each gene sets is: classical n=776 and basal-like n=1,002.

**Moffitt-PuriST**: The PuriST classifier was applied directly on normalized counts using the weights available from the article [13]. The gene sets for basal-like and classical subtypes were extracted from the original NMF [28] assigning to each gene the component for which it has the highest weight. The number of genes in each gene sets is: classical n=879 and basal-like n=692.

**Bailey *et al***.: A centroid classifier was derived using the 1,000 most differentially expressed genes of each subtype versus all others. The number of genes in each gene sets is: Progenitor n=62 and Squamous n=755.

**Puleo *et al***: The centroid classifier from the original study was used. All classes were used, though only the pure tumor classes were identified in PDX. Gene sets were extracted from the ICA definition by selecting all the genes for which the highest weight is found in one of the tumor components. The number of genes in each gene sets is: classical n=830 and basal-like n=733. **Chan-Seng-Yue *et al***.: The gene sets from each NMF tumor component were taken from supplementary Table 4 of the original article. The number of genes in each gene sets is: Classic B n=193, Classic A n=436, Basal B n= 194 and Basal A n=436.

### Unsupervised clustering of PDX

The unsupervised clustering of the 76 PDX transcriptomic profiles was performed as in a previous unsupervised classification of a subset of these PDX [20]. The consensus cluster plus approach was applied on subsets of the most varying genes in the expression matrix using a Pearson distance and aggregating over several linkage methods (average, complete and Ward). Several thresholds of the most varying genes were applied (5% to 25% most variant) and results were aggregated.

### Discovery of Pancreatic Adenocarcinoma Molecular Gradient (PAMG) in PDX

Independent Component analyses were performed on the 50% most variant genes (n=20,434), after gene-wise zero-centering (no unit scaling) and using the JADE algorithm [29]. This resulted in an A matrix of sample projections onto l components defined by a matrix S of gene weights. This matrix decomposition was applied on the 20,026 most variable genes of the 76 PDX with an increasing number of components (2 ≤ *l* ≤ 25). The sample projections of each ICA systems were associated with sample characteristics (e.g. PDX molecular classification and histology classes). Most PDX ICA systems had a component associated with the histological description of the PDX. The ICA system with the component with the highest association with the histological classes (ANOVA, p< 10^−10^) was selected (*l =* 3) and the histological-associated component was identified as the PAMG, which is defined at the sample level by the projection of each sample on the component (from matrix A) and at the gene level by the weights of the selected component (from matrix S).

### Comparing continuum versus dichotomy

From the human compartment of the PDX gene expression, a generalized linear model was fit to each gene in each published signature using either the PAMG or the PurIST classification labels as independent variables. The difference in R2 is reported. As it may be expected that in some cases, the model may better fit to continuous then to discrete independent variables, a background R2-differences was computed on the 25% most variable genes that were not found in any of the published gene set signatures (n=7,393 genes).

### Reproducible identification of the PAMG in other datasets

To reproduce the identification of the PAMG, a blind deconvolution is first applied on the transcriptome profiles and a component with a similar definition than the PDX-based molecular gradient is searched for. Independent Component analyses were performed on the 50% most variant genes/probes of the 309 Affymetrix microarray from Puleo *et al*. (probes n=24,693) and the 269 microarrays from the PDAC ICGC (genes n=23,632). Increasing numbers of components were extracted (2 to 25). For each ICA system of each of these new datasets, the gene weights (matrix S’) were compared to the gene weights of the PDX-based Molecular Gradient ICA system (S), using Spearman’s correlation on the common genes. As shown on supplementary Figure S3, most ICA systems with a sufficient number of components showed at least one component with a high correlation with the PDX-derived PAMG. For each dataset, the system with the highest correlation was selected, and the high correlation component was defined as the dataset-specific PAMG.

### PAMG projection on external cohorts

In order to project a PAMG on a new dataset, the genes found both in the new transcriptome profiles and in the gene weight matrix S of the PAMG ICA system are selected. The cross-product between the Moore-Penrose generalized inverse of the sub-matrix of S and the sub matrix of gene expression with only the common genes is computed.

### Survival analysis

Overall survival was used for survival analysis, using surgery date as the starting of follow-up. Univariate and multivariate Cox proportional hazard regressions were performed using the survival package in R (version 3.6.0). Relative risk plots representing the association between risk of decease and the PAMG were produced by extracting the patient-specific risk of the fitted Cox proportional hazard model, including the 95% confidence interval. Kaplan-Meier curves were generated using t

---

**Supplementary Table I.** Clinical Characteristics of patients included in the study.

| Patient characteristics | n | % |
| --- | --- | --- |
| Total, Median age (66 ) | 76 | 100 |
| Gender |  |  |
| Female | 29 | 38.2 |
| Male | 47 | 61.8 |
| Advancement |  |  |
| Localized | 38 | 50 |
| Locally advanced | 7 | 9.2 |
| Metastatic | 31 | 40.8 |
| Source of material |  |  |
| Curatively resected | 40 | 52.6 |
| EUS-FNA | 25 | 32.9 |
| from Primary tumor | 22 | - |
| from Hepatic meta | 3 | - |
| Surgical biopsies | 11 | 14.5 |
| from Carcinosis | 5 | - |
| from Liver | 4 | - |
| from Lymph node | 1 | - |
| from Primary tumor | 1 | - |
| Survival Status at July 2017 |  |  |
| Alive | 12 | 15.8 |
| Dead | 64 | 84.2 |
| Clinical histol. for Curatively Resected |  |  |
| Well differentiated | 13 | - |
| Moderately differentiated | 15 | - |
| Poor differentiated | 10 | - |
| Missing | 2 | - |

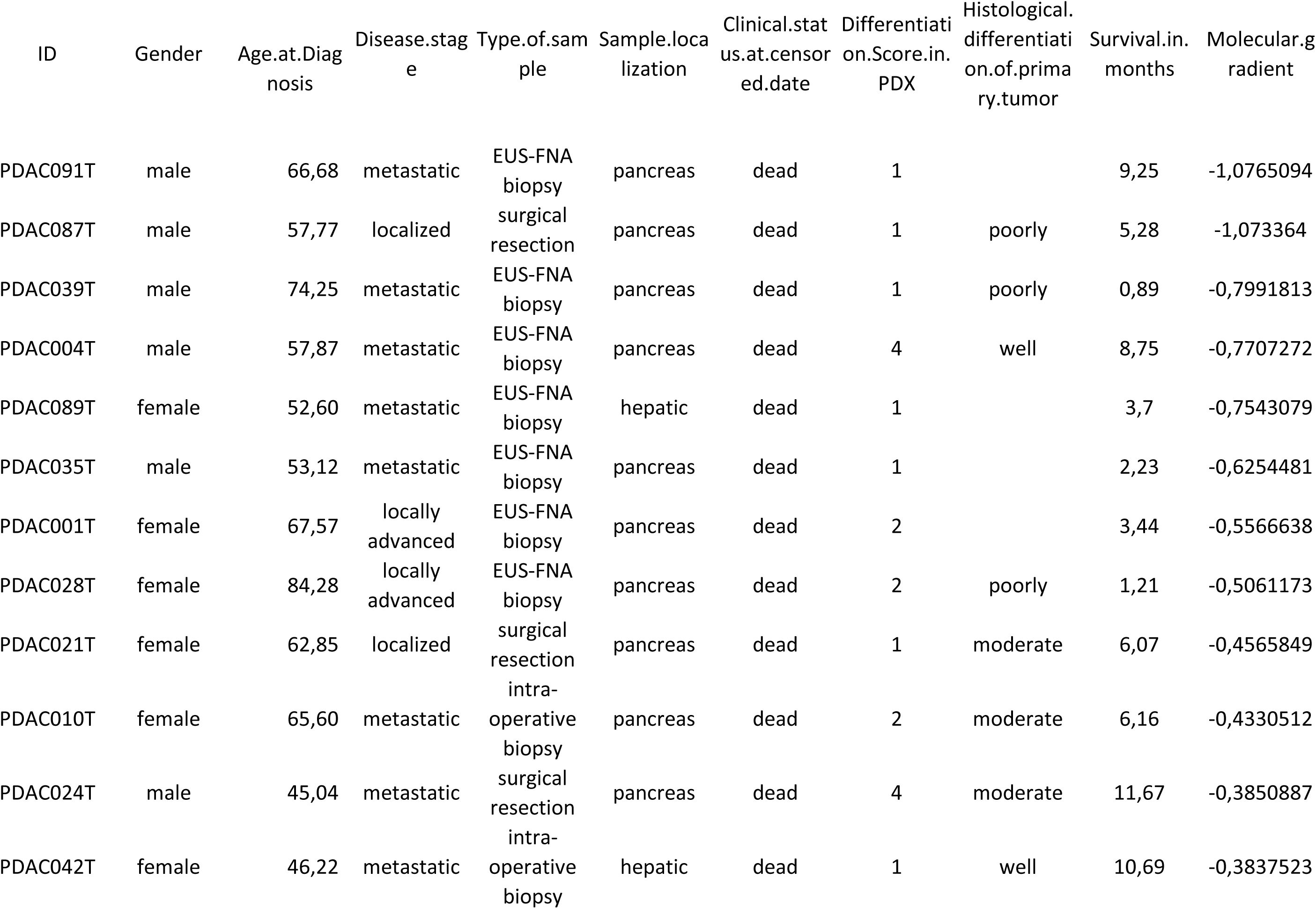
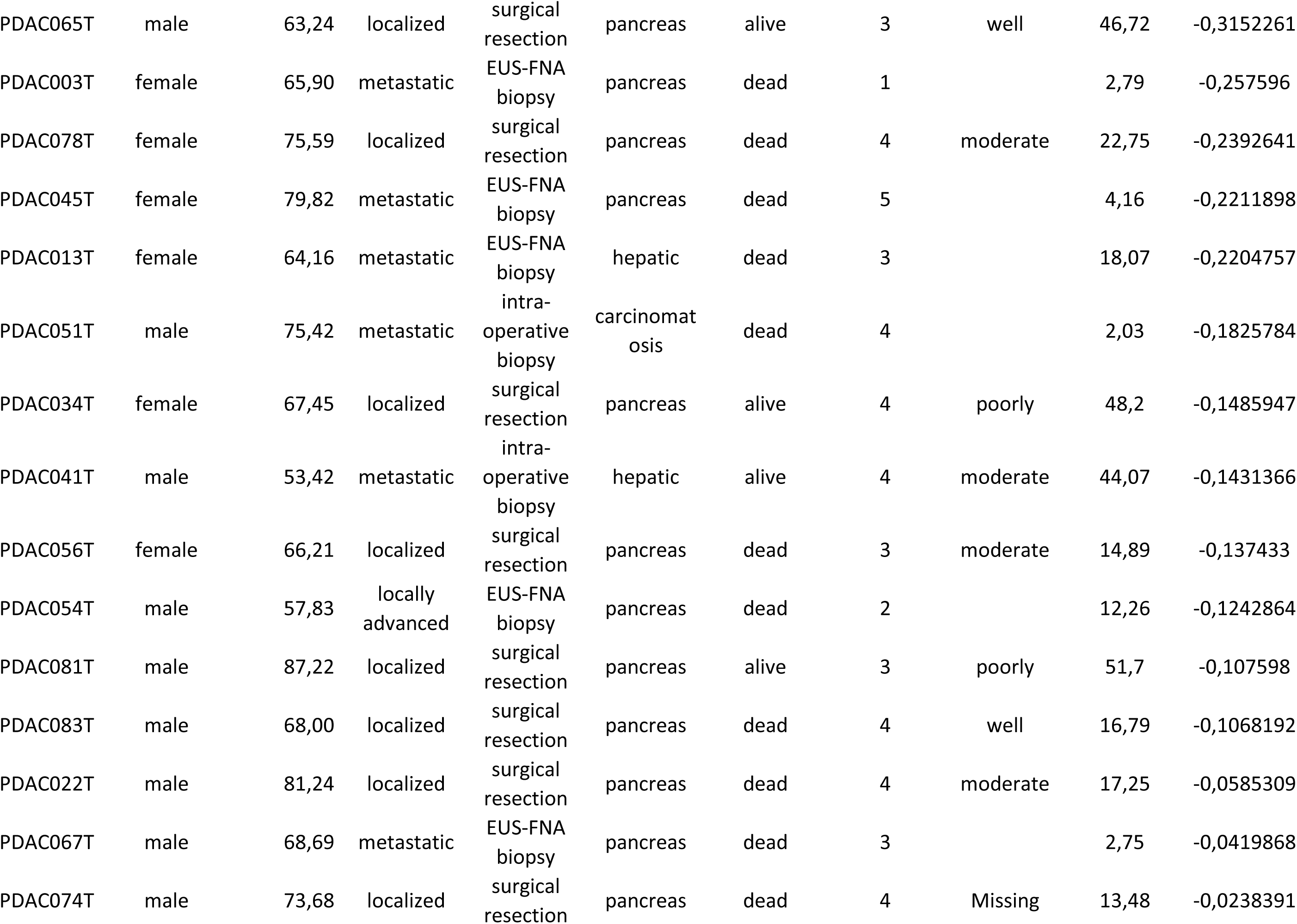
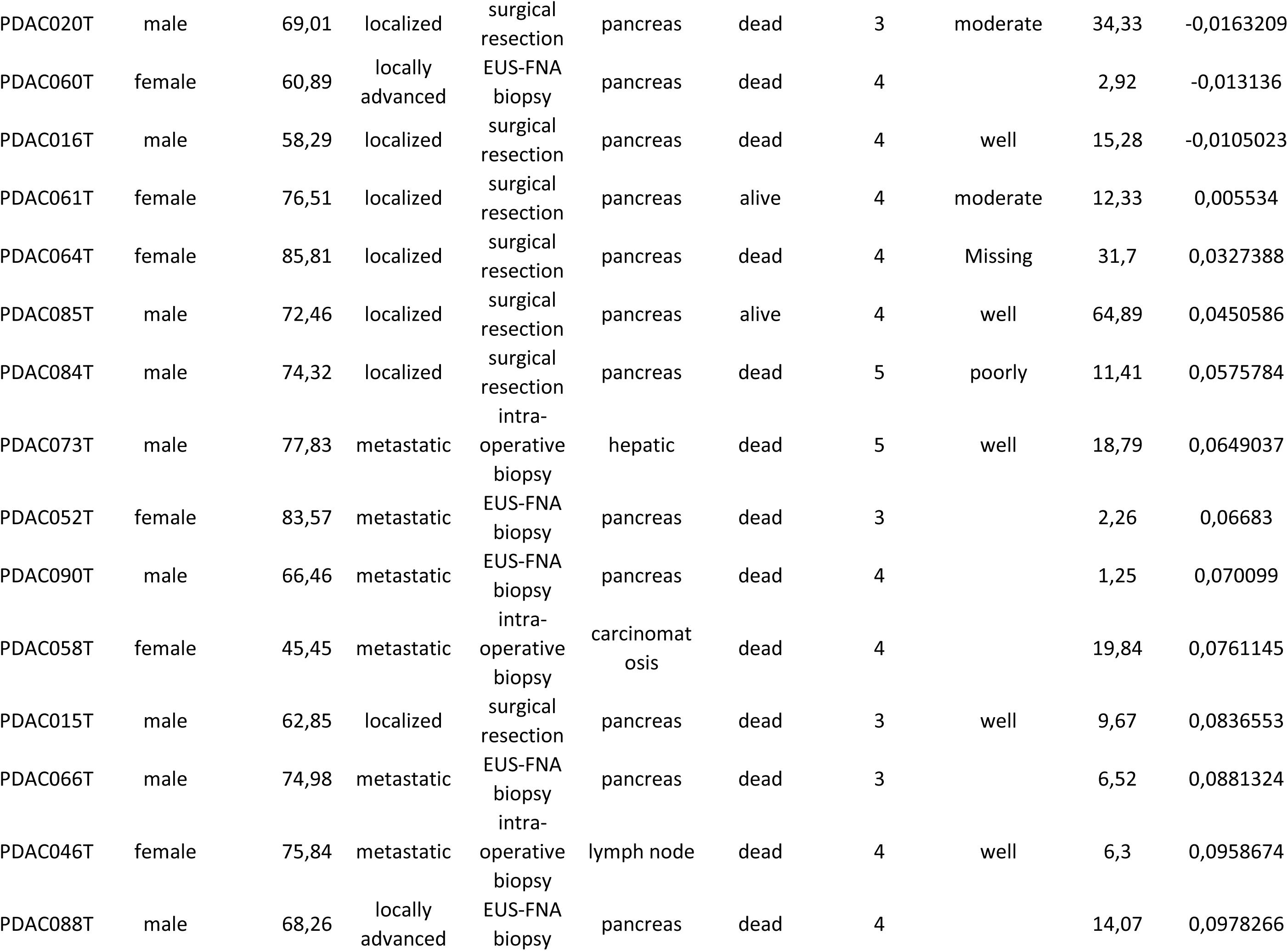
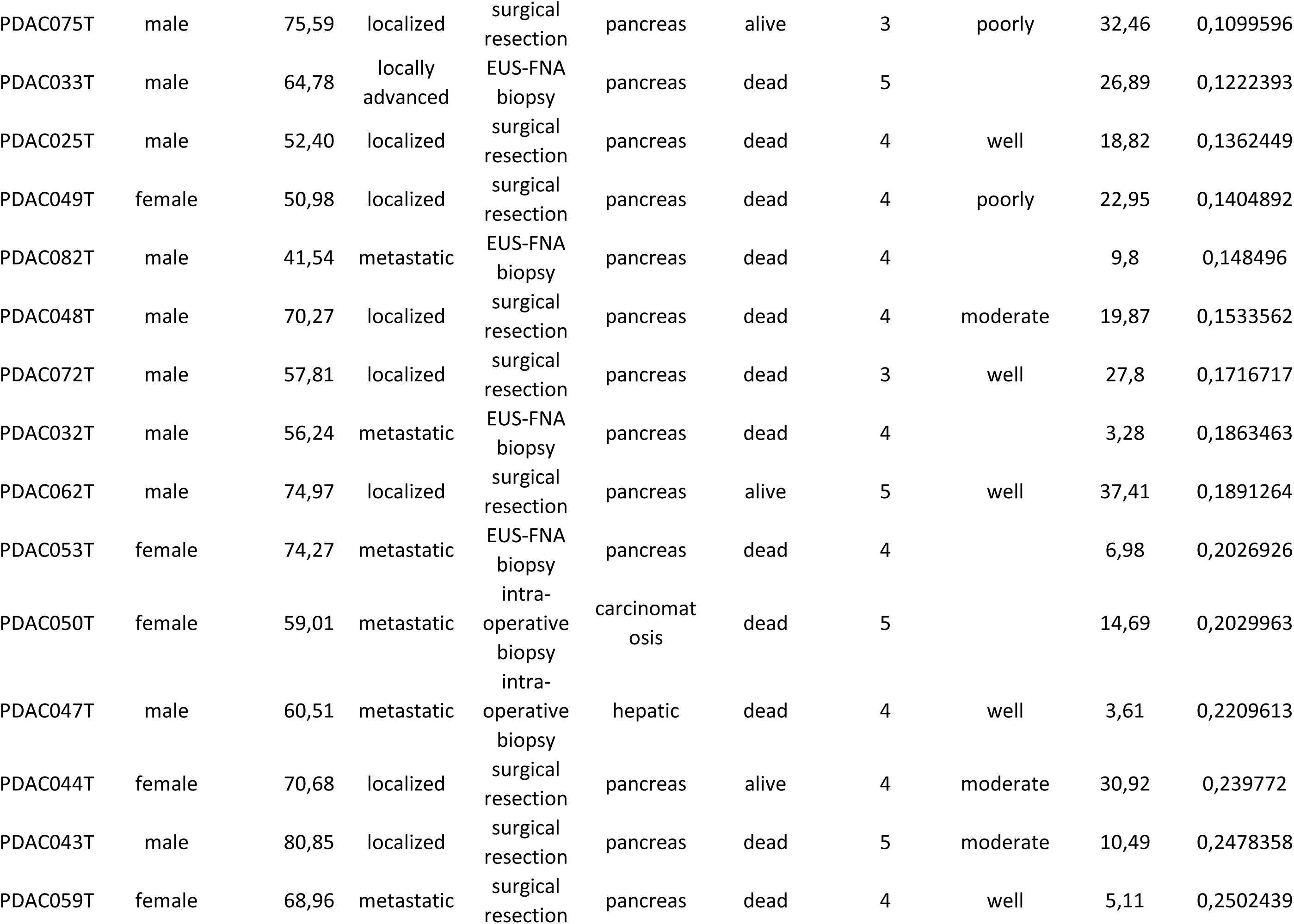
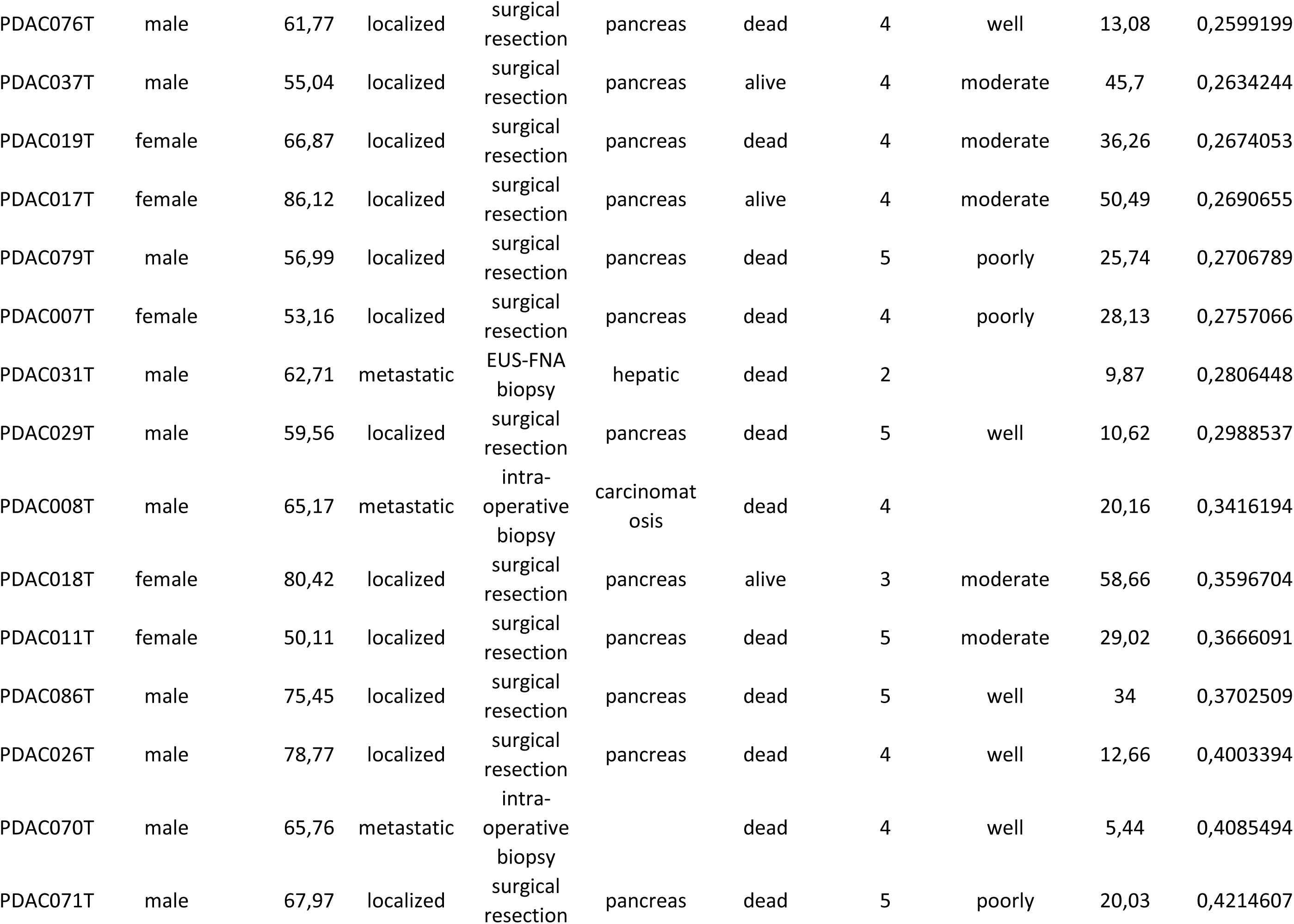
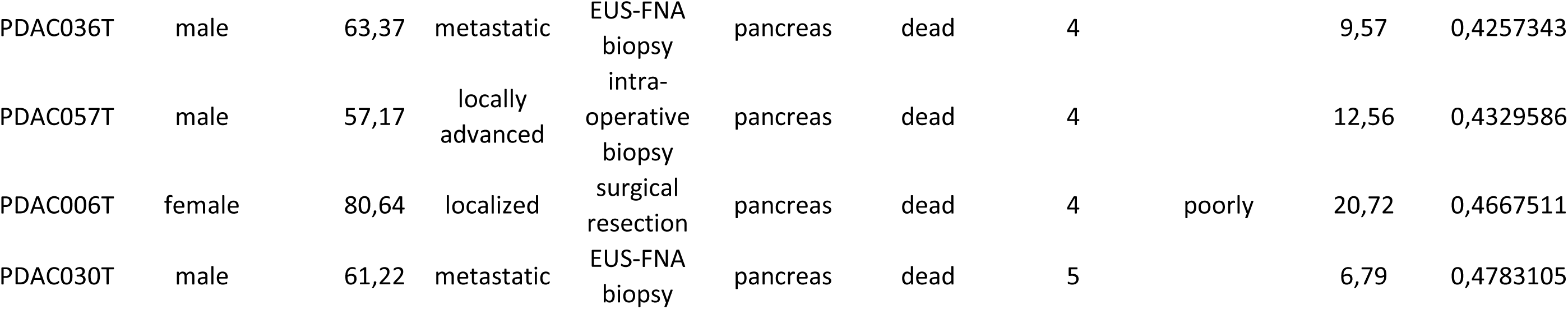

## Bibliography

1. Deramaudt T, Rustgi AK. Mutant KRAS in the initiation of pancreatic cancer. Biochimica et Biophysica Acta (BBA) - Reviews on Cancer 2005; 1756: 97–101.

2. Lenz J, Karasek P, Jarkovsky J et al. Clinicopathological correlations of nestin expression in surgically resectable pancreatic cancer including an analysis of perineural invasion. Journal of gastrointestinal and liver diseases: JGLD 2011; 20: 389–396.

3. Li D, O’Reilly EM. Adjuvant and neoadjuvant systemic therapy for pancreas adenocarcinoma. Semin Oncol 2015; 42:134–143.

4. Hayashi A, Fan J, Chen R et al. A unifying paradigm for transcriptional heterogeneity and squamous features in pancreatic ductal adenocarcinoma. Nature Cancer 2020; 1: 59–74.

5. N Kalimuthu S, Wilson GW, Grant RC et al. Morphological classification of pancreatic ductal adenocarcinoma that predicts molecular subtypes and correlates with clinical outcome. Gut 2019; gutjnl-2019-318217.

6. Nicolle R, Blum Y, Marisa L et al. Pancreatic Adenocarcinoma Therapeutic Targets Revealed by Tumor-Stroma Cross-Talk Analyses in Patient-Derived Xenografts. Cell Reports 2017; 21: 2458–2470.

7. Puleo F, Nicolle R, Blum Y et al. Stratification of Pancreatic Ductal Adenocarcinomas Based on Tumor and Microenvironment Features. Gastroenterology 2018; 155: 1999-2013.e1993.

8. Rashid NU, Peng XL, Jin C et al. Purity Independent Subtyping of Tumors (PurIST), A Clinically Robust, Single-sample Classifier for Tumor Subtyping in Pancreatic Cancer. Clinical Cancer Research 2019; 1078-0432.CCR-1019-1467.

9. Moffitt RA, Marayati R, Flate EL et al. Virtual microdissection identifies distinct tumor- and stroma-specific subtypes of pancreatic ductal adenocarcinoma. Nature Genetics 2015; 47: 1168–1178.

10. Biton A, Bernard-Pierrot I, Lou Y et al. Independent Component Analysis Uncovers the Landscape of the Bladder Tumor Transcriptome and Reveals Insights into Luminal and Basal Subtypes. Cell Reports 2014; 9: 1235–1245.

11. Sompairac N, Nazarov PV, Czerwinska U et al. Independent Component Analysis for Unraveling the Complexity of Cancer Omics Datasets. International Journal of Molecular Sciences 2019; 20: 4414.

12. Lomberk G, Blum Y, Nicolle R et al. Distinct epigenetic landscapes underlie the pathobiology of pancreatic cancer subtypes. Nature Communications 2018; 9.

13. Satelli A, Li S. Vimentin in cancer and its potential as a molecular target for cancer therapy. Cell Mol Life Sci 2011; 68: 3033–3046.

14. Australian Pancreatic Cancer Genome I, Bailey P, Chang DK et al. Genomic analyses identify molecular subtypes of pancreatic cancer. Nature 2016; 531: 47–52.

15. Chan-Seng-Yue M, Kim JC, Wilson GW et al. Transcription phenotypes of pancreatic cancer are driven by genomic events during tumor evolution. Nature Genetics 2020; 52: 231–240.

16. Aung KL, Fischer SE, Denroche RE et al. Genomics-Driven Precision Medicine for Advanced Pancreatic Cancer: Early Results from the COMPASS Trial. Clinical Cancer Research 2018; 24: 1344–1354.

17. Juiz N, Elkaoutari A, Bigonnet M et al. Basal-like and Classical cells coexistence in pancreatic cancer revealed by single cell analysis. bioRxiv 2020; https://doi.org/10.1101/2020.01.07.897454

18. Porter RL, Magnus NKC, Thapar V et al. Epithelial to mesenchymal plasticity and differential response to therapies in pancreatic ductal adenocarcinoma. Proceedings of the National Academy of Sciences 2019; 116: 26835–26845.

19. Mollberg N, Rahbari NN, Koch M et al. Arterial Resection During Pancreatectomy for Pancreatic Cancer: A Systematic Review and Meta-Analysis. Annals of Surgery 2011; 254: 882–893.

20. Bachellier P, Addeo P, Faitot F et al. Pancreatectomy With Arterial Resection for Pancreatic Adenocarcinoma: How Can It Be Done Safely and With Which Outcomes? Annals of Surgery 2018; 1.

21. Bullard JH, Purdom E, Hansen KD, Dudoit S. Evaluation of statistical methods for normalization and differential expression in mRNA-Seq experiments. BMC Bioinformatics 2010; 11.

22. Bournet B, Muscari F, Buscail C et al. KRAS G12D Mutation Subtype Is A Prognostic Factor for Advanced Pancreatic Adenocarcinoma. Clinical and Translational Gastroenterology 2016; 7: e157.

23. Bournet B, Souque A, Senesse P et al. Endoscopic ultrasound-guided fine-needle aspiration biopsy coupled with KRAS mutation assay to distinguish pancreatic cancer from pseudotumoral chronic pancreatitis. Endoscopy 2009; 41: 552–557.

24. Dobin A, Davis CA, Schlesinger F et al. STAR: ultrafast universal RNA-seq aligner. Bioinformatics 2013; 29: 15–21.

25. Liao Y, Smyth GK, Shi W. featureCounts: an efficient general purpose program for assigning sequence reads to genomic features. Bioinformatics 2014; 30: 923–930.

26. Guinney J, Dienstmann R, Wang X et al. The consensus molecular subtypes of colorectal cancer. Nature Medicine 2015; 21: 1350–1356.

27. Kamoun A, de Reyniès A, Allory Y et al. A Consensus Molecular Classification of Muscle-invasive Bladder Cancer. European Urology 2019.

28. Cardoso JF, Souloumiac A. Blind beamforming for non-gaussian signals. IEE Proceedings F Radar and Signal Processing 1993; 140: 362.

